# Discriminative SKP2 interactions with CDK-cyclin complexes support a cyclin A-specific role in p27KIP1 degradation

**DOI:** 10.1101/2020.10.08.329599

**Authors:** Marco Salamina, Bailey C. Montefiore, Mengxi Liu, Daniel J. Wood, Richard Heath, James R. Ault, Lan-Zhen Wang, Svitlana Korolchuk, Arnaud Baslé, Martyna W. Pastok, Judith Reeks, Natalie J. Tatum, Frank Sobott, Stefan T. Arold, Michele Pagano, Martin E.M. Noble, Jane A. Endicott

**Affiliations:** Newcastle University Centre for Cancer, Translational and Clinical Research Institute, Newcastle University, Paul O’Gorman Building, Framlington Place, Newcastle upon Tyne, NE2 4HH, UK; Department of Biochemistry and Molecular Pharmacology, Perlmutter NYU Cancer Center, New York University Grossman School of Medicine, and Howard Hughes Medical Institute, The Alexandria Center of Life Science, East Tower, 450 E, 29thStreet, New York, NY 10016, USA; Astbury Centre for Structural Molecular Biology, School of Molecular and Cellular Biology, University of Leeds, Leeds, LS2 9JT, UK; Biosciences Institute, Newcastle University, Framlington Place, Newcastle upon Tyne, NE2 4HH, UK; Division of Biological and Environmental Sciences and Engineering (BESE), Computational Bioscience Research Center (CBRC), King Abdullah University of Science and Technology (KAUST), Thuwal, 23955-6900, Saudi Arabia; Centre de Biochimie Structurale, CNRS, INSERM, Université de Montpellier, 34090 Montpellier, France

**Keywords:** CDK, cyclin, SKP2, kinase, signaling

## Abstract

The SCF^SKP2^ ubiquitin ligase relieves G1 checkpoint control of CDK-cyclin complexes by promoting p27KIP1 degradation. We describe reconstitution of stable complexes containing SKP1-SKP2 and CDK1-cyclin B or CDK2-cyclin A/E, mediated by the CDK regulatory subunit CKS1. We further show that a direct interaction between a SKP2 N-terminal motif and cyclin A can stabilize SKP1-SKP2-CDK2-cyclin A complexes in the absence of CKS1. We identify the SKP2 binding site on cyclin A and demonstrate the site is not present in cyclin B or cyclin E. This site is distinct from but overlapping with features that mediate binding of p27KIP1 and other G1 cyclin regulators to cyclin A. We propose that the capacity of SKP2 to engage with CDK2-cyclin A by more than one structural mechanism provides a way to fine tune the degradation of p27KIP1 and distinguishes cyclin A from other G1 cyclins to ensure orderly cell cycle progression.

## Introduction

SKP2 (FBXL1) and SKP1 (S-phase kinase-associated proteins 1 and 2) were identified as components of a pentameric cyclin-dependent kinase 2 (CDK2)-cyclin A-cyclin-dependent kinases regulatory subunit 1 (CKS1)-containing complex whose levels were found to be elevated in transformed cell lines compared to their non-tumorigenic counterparts (Goddard et al., 2018; Zhang et al., 1995). SKP2 has since been characterized as a member of a large protein family that shares the F-box motif that binds to SKP1 (Bai et al., 1996; Cenciarelli et al., 1999) and selectively recruits phosphorylated substrates into SCF (SKP1-Cullin1/Cdc53-F-box protein) E3 ubiquitin ligase complexes. These complexes polyubiquitinate their substrates targeting them for degradation by the proteasome (Feldman et al., 1997; Lisztwan et al., 1998; Skowyra et al., 1997; Tsvetkov et al., 1999), reviewed in (Lee and Diehl, 2014; Skaar et al., 2013; Zheng and Shabek, 2017).

The central element of the SKP2 structure (residues 94-140) comprises the F-box that is sandwiched between a larger C-terminal leucine-rich repeat (LRR) domain, and a smaller N-terminal sequence that is predicted to be unstructured. The SKP2 LRR domain and tail recognize phosphorylated substrates that include cyclin-dependent kinase inhibitors (CKIs, including p27KIP1), retinoblastoma-like protein 2 and the transcription factor FOXO1 (reviewed in (Frescas and Pagano, 2008; Heo et al., 2016)). Unusually, SKP2 cooperates with an accessory protein, CKS1, to recognize the phosphorylated T187 residue of p27KIP1 that signals its degradation (Ganoth et al., 2001; Hao et al., 2005; Montagnoli et al., 1999; Sitry et al., 2002; Spruck et al., 2001). The unstructured N-terminal sequence contains a functional D-box near the N-terminus that is recognized by Cdh1 (Bashir et al., 2004; Wei et al., 2004), as well as sequences that interact with cyclin A (Ji et al., 2006) and pRB (Ji et al., 2004). It is also site-specifically phosphorylated (for example by CDK2-cyclin A at S64 (Rodier et al., 2008)), and acetylated, suggesting mechanisms by which protein association might be regulated (Bashir et al., 2010; Cen et al., 2010; Gao et al., 2009; Hao et al., 2005; Inuzuka et al., 2012; Lin et al., 2009; Rodier et al., 2008; Schulman et al., 2000).

The cyclin A binding site on SKP2 has been mapped to four residues within the N-terminal portion of SKP2 (Ji et al., 2006). The character and relative disposition of these residues (L32, L33, S39 and L41 (UniProt entry Q13309 isoform 1 numbering)) suggests a hydrophobic docking sequence that binds to cyclin A. The SKP2 binding site on cyclin A has been proposed to lie within the cyclin A unstructured sequence N-terminal to the two tandem cyclin box folds (Ji et al., 2006). However, SKP2 and p27KIP1 association with CDK2-cyclin A is mutually exclusive suggesting that the SKP2 binding site must overlap with that of p27KIP1 elsewhere on the cyclin A structure (Ji et al., 2006).

Cyclin A is required for DNA replication and, in S-phase, associates with and activates CDK2 (Malumbres, 2014; Morgan, 2007). Cyclin E is an alternative activating partner of CDK2 that is predominantly expressed in late G1 phase. CKIs of the CIP/KIP class (of which p27KIP1 is a member) and selected CDK substrates share a consensus RXL sequence (single letter amino acid code in which X denotes any amino acid) that makes a significant contribution to their binding to CDK-cyclin complexes (Adams et al., 1999). This motif binds to the cyclin recruitment site, a hydrophobic patch on the surface of the N-terminal cyclin box fold (N-CBF) that is conserved in cyclins A, B, D and E (Brown et al., 1999; Russo et al., 1996). The crystal structure of a CDK2-cyclin A-p27KIP1 complex revealed that an N-terminal p27KIP1 fragment both drapes across the surface of cyclin A and integrates itself into the fold of the N-lobe of CDK2 (Russo et al., 1996). p27KIP1 is an intrinsically disordered protein and biophysical characterization of this interaction suggested an induced fit model in which three distinct structural units can be distinguished in the p27KIP1 N-terminal sequence (Lacy et al., 2004; Tsytlonok et al., 2019). The most N-and C-terminal of these units are unstructured in solution and bind to the cyclin A recruitment site and CDK2 N-lobe respectively. The central unit is α-helical and docks onto the surface of cyclin A. RXL-containing CDK substrates are proposed to have more limited interactions with the surface of the CDK-cyclin, docking at the cyclin A recruitment site and thereby effectively increasing local concentration which enhances substrate engagement within the CDK2 catalytic cleft (Cheng et al., 2006; Takeda et al., 2001). Apart from their interactions with CDK partners, no other protein interaction sites have been structurally characterized on the surface of cyclins A, B, D or E that control the cell cycle. In *S. cerevisiae* a conserved “LP” docking site that binds a consensus LXF motif has been identified in Cln1/2 and Ccn1 cyclins adjacent to the recruitment site (Bhaduri et al., 2015).

In this study we identify the features of cyclin A that distinguish it from cyclin E and cyclin B and that mediate formation of a SKP1-SKP2-CDK2-cyclin A-CKS1 pentameric complex. Using purified proteins we show that a stable CDK2-cyclin A-SKP1-SKP2-CKS1 complex can be formed either through CKS1 bridging CDK2 and SKP2 (Hao et al., 2005) or through a direct interaction between the SKP2 N-terminal sequence and a cyclin A SKP2 binding site that we identify in this study. We show that the cyclin A SKP2 and p27KIP1 binding sites have unique and overlapping elements and that SKP2 and p27KIP1 binding to cyclin A is mutually exclusive. We propose a model in which recruitment of non-phosphorylated p27KIP1 to CDK1/2-cyclin A/B/E-SKP1-SKP2-CKS1 complexes is mediated by CKS1. Subsequent phosphorylation of p27KIP1 is promoted by the SKP2 N-terminal cyclin A binding motif that recruits a catalytic CDK-cyclin A to form an octameric complex that promotes phosphorylation of p27KIP1 *in cis*. This SKP2-cyclin A interaction imparts a unique activity to cyclin A to promote p27KIP1 degradation that distinguishes it from cyclins E and B. We hypothesize that the interactions between CDK1 or CDK2 with CKS1, and the interaction between SKP2 and cyclin A act together to determine the efficiency of p27KIP1 T187 phosphorylation and its subsequent SCF^SKP2^-dependent degradation.

## Results

### The cyclin A N-terminal cyclin box fold has a SKP2 binding site

To identify the SKP2 binding site on cyclin A we exploited the knowledge that (i) SKP2 cannot form a pentameric CDK2-cyclin-SKP1-SKP2-CKS1 complex if cyclin E replaces cyclin A (Zhang et al., 1995); (ii) residues in the SKP2 N-terminal sequence prior to the start of the F-box are required for SKP2 to bind to cyclin A (Yam et al., 1999) and (iii) SKP2 and p27KIP1 binding to cyclin A are mutually exclusive (Ji et al., 2006). We first confirmed in a reconstituted system using purified proteins that in the absence of CKS1, cyclin A but not cyclin E can form a complex with CDK2, SKP1 and SKP2 (Figure 1A) and cyclin A but not CDK2 binds directly to SKP1-SKP2 (Supplementary Figure S1A). We next constructed a co-expression system to produce a SKP1-SKP2 complex comprising SKP1 and the first 140 residues of SKP2 (SKP1-SKP2N), encoding the N-terminal regulatory sequence (residues 1-93) and the F-box, (residues 94-140). Using isothermal titration calorimetry (ITC) we confirmed that SKP1-SKP2N binds to CDK2-cyclin A (with an affinity of 298 ± 17 nM (Figure 1B)) but does not bind to CDK2 alone (Table 1 **and** Supplementary Figure S1B). Mapping the sequence conservation between E-cyclins and A-cyclins onto the structure of cyclin A revealed several overlapping surface patches that are highly conserved within but not between the cyclin A and E families. A subset of these sites that are close to, but do not form part of, the cyclin A binding site for p27KIP1 were selected for further study (Table 1, Figure 1 **and** Supplementary Figure S2A).

**Figure 1.**
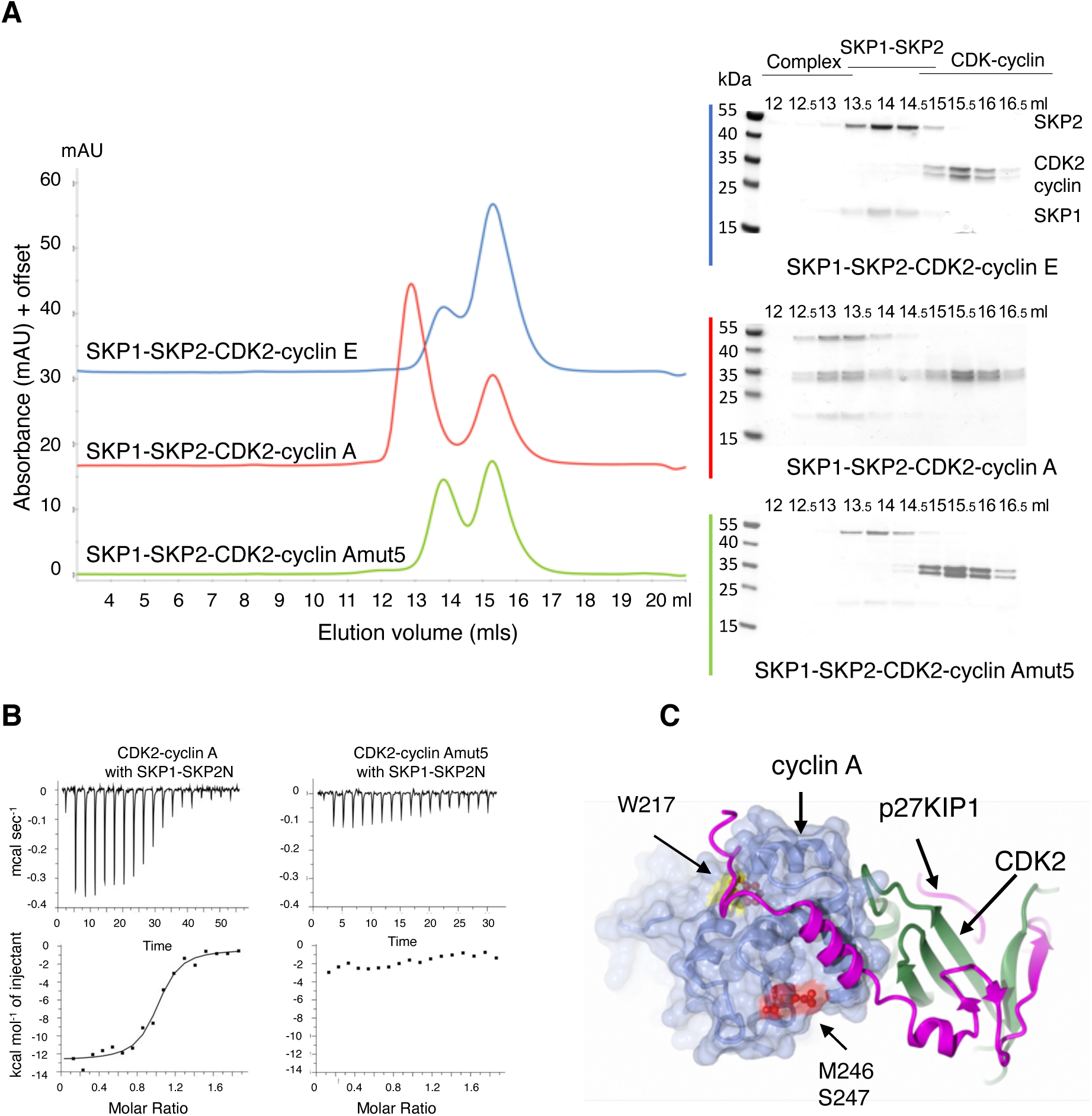
Identification of a SKP2 binding site on cyclin A. (**A**) SKP1-SKP2 forms a stable complex with CDK2-cyclin A, but not with CDK2-cyclin E or CDK2-cyclin Amut5. Chromatograms are to the same scale but have been offset on the y-axis to aid comparison. Accompanying SDS-PAGE analysis. Samples were visualized by Instant Blue staining. Chromatogram is representative of two replicates carried out using independently prepared proteins. Cyclin A and cyclin E residues 174-432 and 96-378 respectively. (**B)** Isothermal titration calorimetry (ITC) thermograms to assess SKP1-SKP2N binding to CDK2-cyclin A and CDK2-cyclin A mutants. ITC thermograms shown are one of two replicates carried out using independently prepared proteins. Data analysis is described in Table 1. (**C)** Locations of cyclin A mutants to identify the cyclin A SKP2 binding site. Structure PDB entry 1JSU.

**Table 1.** Cyclin A mutants display a range of binding affinities to SKP1-SKP2 and p27KIP1. K_d_ values were determined by isothermal titration calorimetry (ITC). ^1^Carried out at 25 °C with a starting concentration of 20 μM CDK2-cyclin A in the cell and 180 μM SKP1-SKP2N in the syringe. ^2^carried out at 25 °C with a starting concentration of 10 μM CDK2-cyclin A in the cell and 120 μM SKP1-SKP2N or p27KIP1MS in the syringe. ^3^not determined, ^4^not detectable. Experiments were carried out using a Microcal ITC 200. ^5^Kd is the derived average of two independent biological replicates and SD of the population. ^6^Value derived from one measurement where the error represents the fitting error directly derived from the thermogram.

| Cyclin A | Residues mutated | Residue changes | SKP1-SKP2N<br>K<sub>d</sub> ± (nM) | p27KIP1MS<br>K<sub>d</sub> ± (nM) |
| --- | --- | --- | --- | --- |
| <sup>1</sup> Wild-type | none | none | 298 ± 17<sup>5</sup> | 350 ± 36<sup>5</sup> |
| <sup>1</sup> mut2 | 284-289 | DTYTKK to GACSGD | 340 ± 60<sup>6</sup> | -.<sup>3</sup> |
| <sup>1</sup> mut3 | 290-299 | QVLRMEHLVL to<br>EILTMELMIM | 658 ± 101<sup>6</sup> | - |
| <sup>1</sup> mut5 | 246-247 | MS to QEN | ND<sup>4, 5</sup> | 176 ± 1.5<sup>5</sup> |
| <sup>1</sup> mut6 | 299-305 | LKVLTFD to MKALKWR | 1690 ± 340<sup>6</sup> | - |
| <sup>1</sup> mut7 | 244-247 | SSM-S to ATQEN | ND<sup>6</sup> | - |
| <sup>1</sup> global | 244-247,<br>284-305 | S244A, S245T, M246Q,<br>S247E+N, D284G, T285A,<br>Y286C, T287S, K288G,<br>K289D, Q290E, V291I,<br>R293T, H296L, L297M,<br>V298I, L299M, V301A,<br>T303K, F304W, D305R | ND<sup>6</sup> | - |
| <sup>2</sup> W217 | 217 | W217K | ND<sup>5</sup> | ND<sup>5</sup> |
| <sup>2</sup> M210 | 210 | M210D | 894 ± 120<sup>6</sup> | 401 ± 177<sup>6</sup> |
| <sup>2</sup> I213 | 213 | I213D | 50.7 ± 11.0<sup>5</sup> | 1800 ± 160<sup>5</sup> |
| <sup>2</sup> R293 | 293 | R293E | 728 ± 282<sup>5</sup> | ND<sup>5</sup> |
| <sup>2</sup> L297 | 297 | L297K | ND<sup>5</sup> | ND<sup>5</sup> |

To test these potential SKP2 binding sites we again used ITC to compare the binding of SKP1-SKP2N to the authentic cyclin A and cyclin A mutant proteins (Table 1 **and** Supplementary Figure S2B). Where all the selected sequence loci in cyclin A were mutated to replicate the sequence of cyclin E (generating a variant termed cyclin Aglobal), binding to SKP1-SKP2N was completely abrogated. Individually mutating certain subsets of these sequence loci (284-289 to generate cyclin Amut2, 290-299 to generate cyclin Amut3 or 299-305 to generate cyclin Amut6) had no discernible or relatively little impact on SKP2-cyclin A association. By contrast, changing the cyclin A sequence at residues 244-247 (SSMS, Supplementary Figure S2B) or at 246-247 (MS, Figure 1B, C) to the equivalent cyclin E sequences (ATQEN or QEN respectively, generating the respective variants cyclin Amut7 and cyclin Amut5 had a profound effect on the interaction. To verify the ITC results we used SEC to confirm that full-length SKP2-SKP1 does not form a complex with CDK2-cyclin Amut5 (Figure 1A). Finally, to confirm the structural and functional integrity of cyclin Amut 5 we prepared the CDK2-cyclin Amut5 complex and both determined its crystal structure (Supplementary Figure S2C, Supplementary Table S1, PDB entry 6SG4) and compared its activity with CDK2-cyclin A towards a model RXL-containing peptide substrate derived from p107 (Brown et al., 2015) (Supplementary Figure S2D **and** Supplementary Table S2). The structure shows that the insertion has not affected the overall fold and the *k*_cat_/K_m_ values for each complex towards this substrate were comparable.

### Cyclin A has distinct but overlapping SKP2 and p27KIP1 binding sites

This site which we have termed the “SSMS site”, identified by cyclin Amut5, represents a novel site of cyclin A protein interaction. It is distant from the cyclin recruitment site but is close to residues that interact with p27KIP1 (Figure 1C). Given the locations of the SSMS site, the RXL recruitment site and the extended p27KIP1 α-helical cyclin A binding sequence we hypothesized that the extended SKP2 binding site might overlap with either the p27KIP1 RXL motif or α-helical cyclin A binding sites. (Respectively also referred to as p27KIP1 sequence “D1” (residues 27-37) and “linker helix” (residues 38-58) (Huang et al., 2015)).

To test this model, we first confirmed using surface plasmon resonance (SPR) that the binding of cyclin A to p27KIP1 was not affected by the mut5 mutation (Supplementary Figure S3A). GST-p27KIP1_1-106_ (p27KIP1M) was immobilized on the SPR chip surface and CDK2-cyclin A and CDK2-cyclin Amut5 were flowed over as analytes. The K_d_ values determined by this method were not significantly different, being respectively 15.5 ± 0.88 nM and 25.00 ± 0.82 nM (p > 0.05).

We next prepared a series of cyclin A variants, mutated at residues that interact with p27KIP1 or are located close to the recruitment or SSMS sites, and tested their ability to bind to p27KIP1 and to SKP1-SKP2 using ITC (Figure 2A, Table 1 **and** Supplementary Figure S3B). For this assay we again used the SKP1-SKP2N construct and, in order to assess only the cyclin A binding properties of p27KIP1, we used a truncated fragment of p27KIP1 encompassing residues 23 to 51 (p27KIP1MS) that includes the RXL motif and does not contact CDK2. Measured under the ITC conditions, p27KIP1MS binds to CDK2-cyclin A with an affinity of 350 ± 36 nM (Table 1).Variants of cyclin A were prepared to probe the contribution to protein binding made by the recruitment site (variant cyclin A_W217K), by an adjacent hydrophobic pocket that binds to F33 of p27KIP1 (variant cyclin A_I213D), by a residue from helix α5 that does not directly contact p27KIP1 (variant cyclin A_M210D), and by the cyclin A surface groove that binds the p27KIP1 linker helix (variants cyclin A_R293E and cyclin A_L297K). Cyclin A_R293E was engineered because R293 interacts with p27KIP1 E46, whereas cyclin A_L297K was chosen because L297 contributes to a hydrophobic pocket that accepts p27KIP1 residues L45 and C49 and the comparable residue is a lysine in cyclin B. ((Russo et al., 1996), Figure 2A). As expected, cyclin Amut5 bound tightly to p27KIP1 (Supplementary Figure S3B, Table 1) but cyclin A mutants W217K, I213D, R293E (Figure 2A and Table 1) and L297K (Supplementary Figure S3B and Table 1) showed reduced binding to p27KIP1 (significance calculated at p < 0.05). The introduction of the W217K mutation also disrupted binding to SKP1-SKP2N (Figure 2A) suggesting that SKP2 binding also depends on the cyclin partner having an intact recruitment site, despite SKP2 not containing a canonical RXL motif. We confirmed W217K mutation does not affect the overall integrity of the cyclin A fold by differential scanning fluorimetry (Supplementary Figure S3C). CDK2-cyclin A, CDK2-cyclin Amut5 and CDK2-cyclin AW217K have comparable T_m_ mean values of 48.75(±0.22) °C, 47.05 (±0.23) °C and 47.68(±0.42) °C respectively.

**Figure 2.**
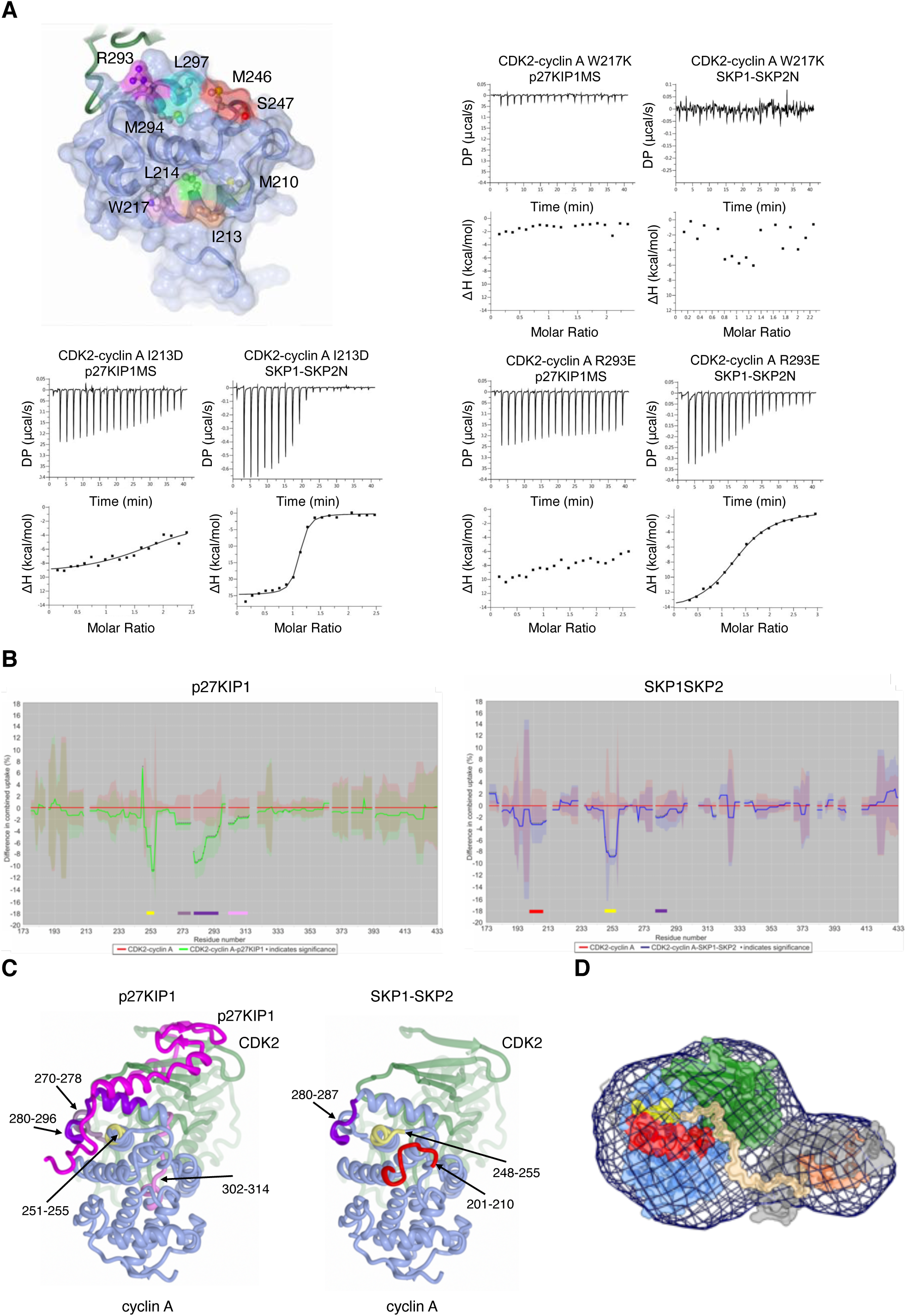
Defining the SKP2 binding site on cyclin A. (**A**) Cyclin A mutants distinguish SKP2 and p27KIP1 binding sites on cyclin A. Isothermal titration calorimetry (ITC) thermograms for the binding of p27KIP1MS and SKP1-SKP2N to cyclin A mutants. ITC thermograms shown are of one of two replicates carried out using independently prepared proteins. Data analysis is described in Table 1. (**B**) Hydrogen-deuterium exchange mass spectrometry distinguishes p27KIP1 and SKP2 binding to cyclin A. Differences in the combined relative fractional uptake of CDK2-cyclin A alone and in the presence of p27KIP1M (LHS) or SKP1-Δ20SKP2N (RHS) is visualized using the PAVED algorithm (Cornwell et al., 2018) and plotted against cyclin A residue number. Uptake plots shown were determined after 2.0 minutes deuterium incubation. Significant differences are marked by asterisks and the colored bars highlight areas discussed in the text. (**C**) Mapping of the HDX-MS footprints onto the structure of CDK2-cyclin A-p27KIP1 (PDB entry 1JSU). Coloring as in (B). (**D**) SAXS-derived model for the CDK2-cyclin A-SKP1-SKP2N complex. CDK2 and cyclin A are colored as in Figure 1, the SKP2 F-box and SKP1 are colored salmon and grey respectively. The cyclin A SSMS and p27KIP1 binding sites are identified in red and lemon respectively and the SKP2 N-terminal sequence is modelled in khaki. Mesh represents the volume-adjusted (damfilt.pdb) converted into a map using molmap and visualized in ChimeraX (Goddard et al., 2018).

Mutant M210D at the start of the MRAIL sequence had little effect on the binding of either p27KIP1 or SKP2 (Supplementary Figure S3B). This residue does not contact p27KIP1 in the CDK2-cyclin A-p27KIP1 structure and this result suggests that it lies distant from the SKP2 binding site as well. Intriguingly, introduction of the I213D mutation reduced affinity for p27KIP1 *circa* 5-fold, but enhanced affinity for SKP2 *circa* 5-fold (Figure 2A **and** Table 1). Cyclin A mutations in α5 also distinguish p27KIP1 and SKP2 binding, suggesting that p27KIP1 and SKP2 do not share common interactions with this region. Binding of SKP2 was severely compromised by the introduction of the L297K mutation into cyclin A, but we did not measure a significant effect for the R293E mutation that ablates p27KIP1MS binding (p > 0.05).

We next characterised CDK2-cyclin A, CDK2-cyclin A-p27KIP1 and CDK2-cyclin A-SKP2-SKP2 complexes by hydrogen-deuterium exchange mass spectrometry (HDX-MS) to distinguish p27KIP1 and SKP2 binding to cyclin A by an orthogonal technique, and to explore the extent of the cyclin A surface affected by SKP2 binding (Figure 2B, Supplementary Figure S4). For these experiments, p27KIP1M (residues 1-106) and Δ20SKP2N (residues 20-140) constructs were employed. We determined the comparative uptake profiles for CDK-cyclin A on its own, or in complex with either p27KIP1M or SKP1-Δ20SKP2N. Determination of deuterium uptake was made at the peptide level (following online pepsin digestion of the proteins) for deuterium incubation times of 30 s, 1 min, 2 min, 10 min and 60 min. Following data processing, differential exchange between the conditions was identified using PAVED software (Cornwell et al., 2018) to determine the highlighted statistically significant differences in deuterium uptake (Figure 2B, Supplementary Figure S4). The significant areas of modulated exchange on cyclin A are mainly focussed around five regions, however the exchange ‘fingerprint’ is unique in either case. In the presence of p27KIP1M four areas show increased protection (Figure 2B, C, yellow, lilac, purple and pink) corresponding to positions 251-255, 270-278, 280-296 and 302-314 respectively. As expected, these sequences map closely to the p27KIP1 binding site ((PDB 1JSU)) and to residues R293 and L297 mutation of which affects p27KIP1 binding (Table 1). In contrast in the CDK2-cyclin A-SKP1-Δ20SKP2N complex, cyclin A peptides show significant protection at positions 201-210 (red), 248-255 (yellow and includes the SSMS sequence), and to a smaller extent 280-287 (purple) (Figure 2B**, C**). Mutation of cyclin A residues I213, W217 and L297 also affect p27KIP1 and SKP2 binding to cyclin A (Table 1). However, fragments 213-218 and 295-299 that respectively include these residues were already highly protected even in the absence of p27KIP1M and Δ20SKP2N SKP2 (Supplementary Figure S4), and did not undergo significant exchange in either the CDK2-cyclin A-p27KIP1 or CDK2-cyclin A-Δ20SKP2N complexes over the time course of the experiment.

Taken together the results of the ITC and HDX-MS experiments suggest a model in which p27KIP1 and SKP2 engage with the cyclin A RXL site and make related but distinct contacts with the N-terminal end of the MRAIL motif. However, their paths subsequently diverge: whereas p27KIP1 interacts with the start of α5 to traverse the “top” of the cyclin A fold, SKP2 does not. Both protein binding sites then converge again at the C-terminal end of cyclin A α5 as defined by their shared sensitivity to the mutation L297K (Figure 2C).

### An extended SKP2 sequence binds to cyclin A

Previous work has shown that SKP2 residues L32, L33, S39 and L41 interact with cyclin A (Ji et al., 2006). To confirm the importance of these residues and assess the extent of the SKP2 sequence that binds to cyclin A we prepared a series of authentic and mutant SKP1-SKP2N constructs and short SKP2 peptides, and assessed their binding to CDK2-cyclin A by ITC (Table 2, Supplementary Figure S5). Compared to binding of the authentic sequence, truncating SKP2N to start at F20 had little effect on the affinity (Supplementary Figure S5A). SKP1-SKP2N_4A (with mutations L32A, L33A, S39A and L41A) showed *circa* 15-fold decrease in binding to CDK2-cyclin A suggesting that other SKP2 residues contribute ((Supplementary Figure S5B) and compare with Supplementary Figure S3B). Although the SKP2 N-terminal sequence contains a number of residues that are highly conserved, two tryptophan residues at positions W22 and W24 are prominent. Binding of SKP2 to cyclin A was not detectable when these residues were also mutated to alanine to generate SKP1-SKP2N_6A (mutations SKP2_4A and W22A and W24A) (Supplementary Figure S5C). Thus, the SKP2 sequence between residues F20 and E45 contribute substantially to cyclin A binding and we propose that this sequence constitutes a SKP2 cyclin A interacting motif (CAIM).

**Table 2.** SKP2 mutants display a range of binding affinities to cyclin A. K_d_ values were determined by isothermal titration calorimetry. ^1^Carried out at 25 °C with a starting concentration of 10 μM CDK2-cyclin A in the cell and 120μM SKP1-SKP2N in the syringe. ^2^not detectable. Experiments were carried out using a Malvern Microcal PEAQ-ITC. K_d_ is the derived average of two independent biological replicates and SD of the population.

| Skp2 | Residue changes | Cyclin A<br>$K_d \pm$ (nM)<sup>1</sup> |
| --- | --- | --- |
| Wild-type | None | 298 ± 17 |
| SKP2N_4A | L32A L33A S39A L41A | 5290 ± 215 |
| SKP2N_6A | W22A W24A L32A L33A S39A L41A | ND<sup>2</sup> |
| SKP2N_Δ20 | N-terminal 20 residues deleted | 507 ± 295 |

While an intact CAIM is required for SKP2 to bind to cyclin A, we were not able to detect stable binding of isolated CAIM peptides to cyclin A using ITC (Supplementary Figure S5D-F), suggesting that the interaction is dependent on the context in which the CAIM is presented. To characterize this context-dependent interaction we analyzed a solution of CDK2-cyclin A and SKP1-SKP2N by small angle X-ray scattering (SAXS) (Figure 2D, Supplementary Figure S6, Supplementary Table S3). This analysis suggests a stable, fairly compact and rigid assembly, suggesting that the SKP1-SKP2 module “scaffolds” the CAIM as it binds to cyclin A, thereby rigidifying the resulting SKP1-SKP2N-CDK2-cyclin A complex.

### SKP2 binds to CDK2-cyclin A-CKS1 through two distinct interactions

A tight (nM) interaction between CKS1 and the C-terminal sequence of SKP2 helps stabilize the pentameric CDK2-cyclin A-CKS1-SKP1-SKP2 complex (Hao et al., 2005). To assess the relative contributions of the CKS1-SKP2 and cyclin A-SKP2 interactions to the stability of the CDK2-cyclin A-CKS1-SKP1-SKP2 pentameric complex, two sets of mutations were made to block each interaction and the resulting proteins were analyzed by SEC. To compromise the binding of SKP2 to CKS1, a SKP2 variant was constructed in which F393 was mutated to a glycine-serine pair (SKP2_C2, Supplementary Figure S7A), while cyclin Amut 5 was used to compromise the binding of cyclin A to the CAIM. By SEC, a stable pentameric complex forms when either the SKP2-cyclin A (green trace) or CKS1-SKP2 (blue trace) interaction is intact, but in the absence of both sites the proteins migrate as CDK2-cyclin A-CKS1 and SKP1-SKP2 sub-complexes (orange trace, Figure 3A). This analysis demonstrates that either site is sufficient to stabilize the pentameric complex in cell free systems. Incubating SKP1-SKP2 with an excess of CDK2-cyclin A-CKS1 did not lead to the formation of a larger complex indicating that a (CDK2-cyclin A-CKS1)_2_-SKP1-SKP2 complex is not abundant under these conditions (Figure 3A, magenta trace). However, we cannot exclude that such a complex might form dependent on effective local concentrations of CDK2-cyclin A. Using SKP1-SKP2, CDK2-cyclin E-CKS1 and CDK1-cyclin B-CKS1 we could also demonstrate that the CDK2-CKS1 or CDK1-CKS1 interface is sufficient to maintain a stable pentameric complex in the absence of a cyclin A-SKP2 interaction (Supplementary Figure S7B). These results demonstrate that in the absence of CKS1, only CDK-cyclin A (and not -cyclin B or -cyclin E) can form a stable complex with SKP1-SKP2. However, if CKS1 is present, then it can bind to either CDK1 or CDK2 and SKP2 to “glue” the CDK-cyclin and SKP1-SKP2 binary complexes together. Furthermore, SEC analysis shows that p27KIP1-CDK2-cyclin A forms a stable complex with SKP1-SKP2 only in the presence of CKS1 (Supplementary Figure S7C). Taken together these observations suggest a model in which the recruitment of p27KIP1 into a complex with SKP1 and SKP2 is dependent on the presence of CKS1 and is independent of the identity of its CDK-cyclin partners.

**Figure 3.**
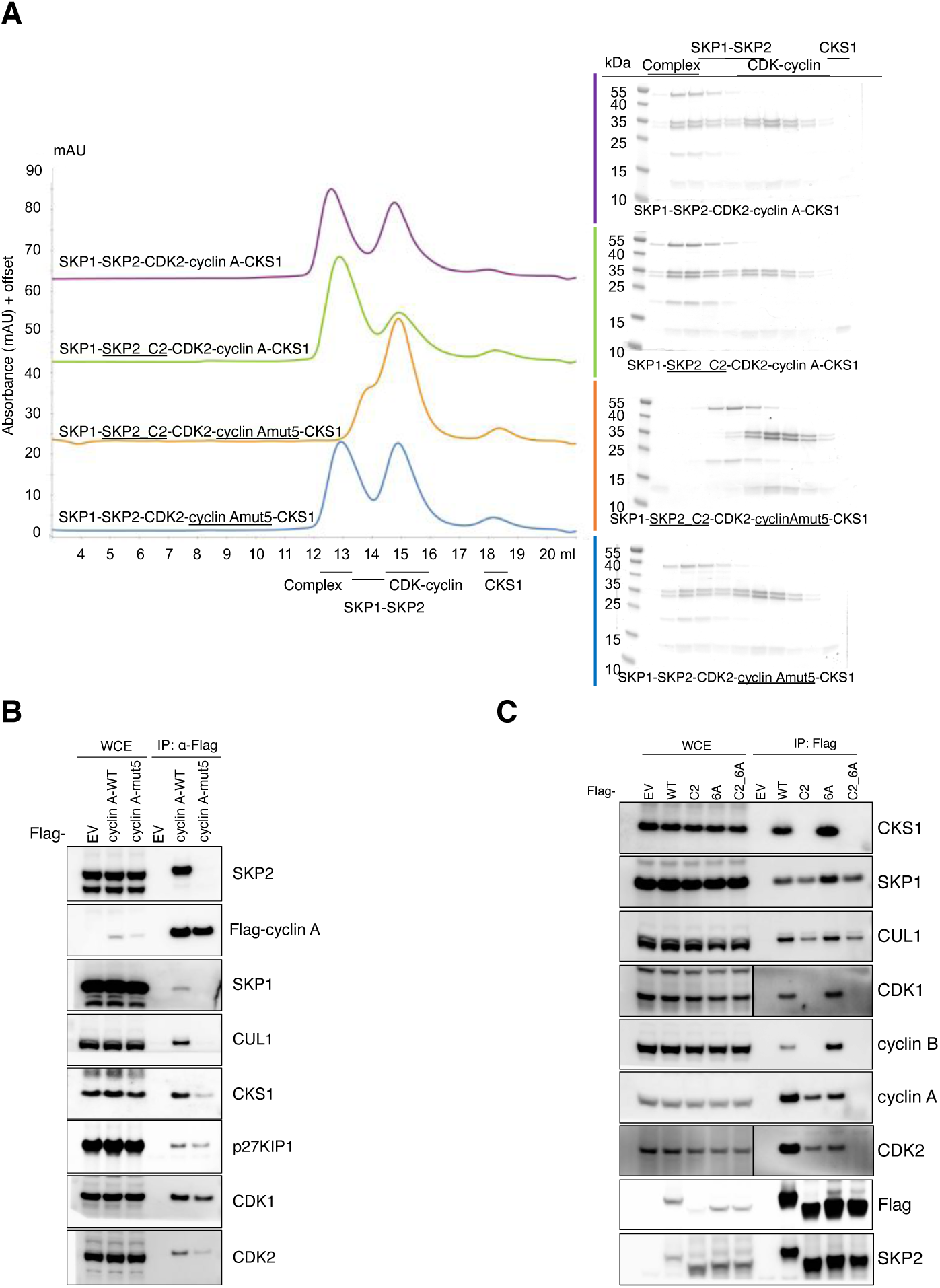
Characterization of the CDK2-cyclin A-SKP1-SKP2-CKS1 complex. (**A**) Size exclusion chromatography to assess the contribution of the SKP2-cyclin A and SKP2-CKS1 interfaces to generation of a stable pentameric complex. Accompanying SDS-PAGE analyses are provided in the color-matched boxes. Proteins were visualized using InstantBlue. Chromatograms are to the same scale but have been offset on the y-axis to aid comparison. Each chromatogram is representative of 2 replicates carried out using independently prepared proteins. (**B**) HEK293T cells were transfected with either an empty vector (EV) or FLAG-tagged cyclin A_WT or cyclin A_mut5. Twenty-four hours post-transfection, cells were collected for immunoprecipitation (IP) and immunoblotting. WCE, whole-cell extract. (**C**) HEK293T cells were transfected with either an empty vector (EV) or FLAG-tagged SKP2_6A, SKP2_C2, or SKP2_C2_6A. The experiment was performed as described in Figure 3B.

### The cyclin A SSMS site promotes cyclin A-SKP2 interaction *in cellulo*

To evaluate the functional significance of the cyclin A SSMS site *in cellulo*, we first transfected full-length cyclin A, and full-length cyclin Amut5 into HEK293T cells (Figure 3B). Immunoprecipitation exploiting the FLAG tag followed by western blotting revealed that whereas a robust interaction can be detected between authentic cyclin A and SKP2, introduction of the mut5 mutation abolished binding. Moreover, compared to wild type cyclin A, cyclin Amut5 has a reduced interaction with SKP1, CKS1 and CDK2. These results demonstrate that the SSMS site makes a significant contribution to the stability of the pentameric CDK2-cyclin A-CKS1-SKP1-SKP2 complex in a cellular setting. This site is not present in cyclin E and helps to distinguish the abilities of these two cyclins to form stable complexes with SKP2.

To further assess the direct interaction between SKP2 and cyclin A and the non-direct interaction mediated by CKS1, N-terminally FLAG-tagged full-length authentic SKP2, SKP2_6A, SKP2_C2 and SKP2_6A_C2 were expressed in HEK293T cells to test their ability to bind to cyclin A (Figure 3C). The wild type SKP2 interacts robustly, but mutation of either the cyclin A-SKP2 (SKP2_6A mutant) or the CKS1-SKP2 interface (SKP2_C2 mutant) significantly decreases association. As expected, the cyclin A-SKP2 interaction is totally abolished when both binding sites are inactivated. Taken together these results support a model in which either the cyclin A-SKP2 or CKS1-SKP2 interaction can mediate complex formation, but that loss of both interaction sites abrogates binding.

### The cyclin A-SKP2 interaction recruits CDK2-cyclin A to promote p27KIP1 phosphorylation

p27KIP1 can inhibit CDK1 and CDK2 bound to cyclins A and B, or A and E respectively. Relief of p27KIP1 inhibition and cell cycle progression results from CDK-dependent p27KIP1 phosphorylation at T187 and its subsequent targeting to the SCF^SKP2^ E3 ubiquitin ligase. Our finding that p27KIP1 and the CAIM of SKP2 share overlapping footprints on cyclin A suggests that binding of p27KIP1 to a pentameric CDK1/2-cyclin A-CKS1-SKP1-SKP2 complex would displace the CAIM from its binding site on cyclin A, making it available to form alternative interactions. Because neither cyclin B or cyclin E possess a site that binds the CAIM, we would similarly expect the CAIM to be available in the pentameric CDK1/2-cyclin B-CKS1-SKP1-SKP2 and CDK1/2-cyclin E-CKS1-SKP1-SKP2 complexes. A potential role for the CAIM in these complexes might be to recruit a further complex of CDK1/2-cyclin A (but not CDK1/2-cyclin B or CDK1/2-cyclin E). This additional copy might play a catalytic role in phosphorylating the C-terminus of p27KIP1.

To test this model, we carried out SEC to evaluate whether a stable octameric complex might form when SBP-tagged CDK2-cyclinA (magenta trace, Figure 4A) is incubated with CDK2-cyclin A-CKS1-p27KIP1-SKP1-SKP2 (blue trace). Our results confirm the formation of such an octameric complex (orange trace), and further demonstrate that complex stability is dependent on the integrity of the SSMS site: no octameric complex is formed when SBP-tagged CDK2-cyclin Amut5 is incubated with CDK2-cyclin A-CKS1-p27KIP1-SKP1-SKP2 (green trace). We can also infer that the recruitment of the second CDK2-cyclin A module is dependent on p27KIP1 being present: SEC analysis of CDK2-cyclin A-SKP1-SKP2-CKS1 in the presence of excess CDK2-cyclin A does not indicate formation of (CDK2-cyclin A)_2_-SKP1-SKP2-CKS1 complex (Figure 3A, magenta trace).

**Figure 4.**
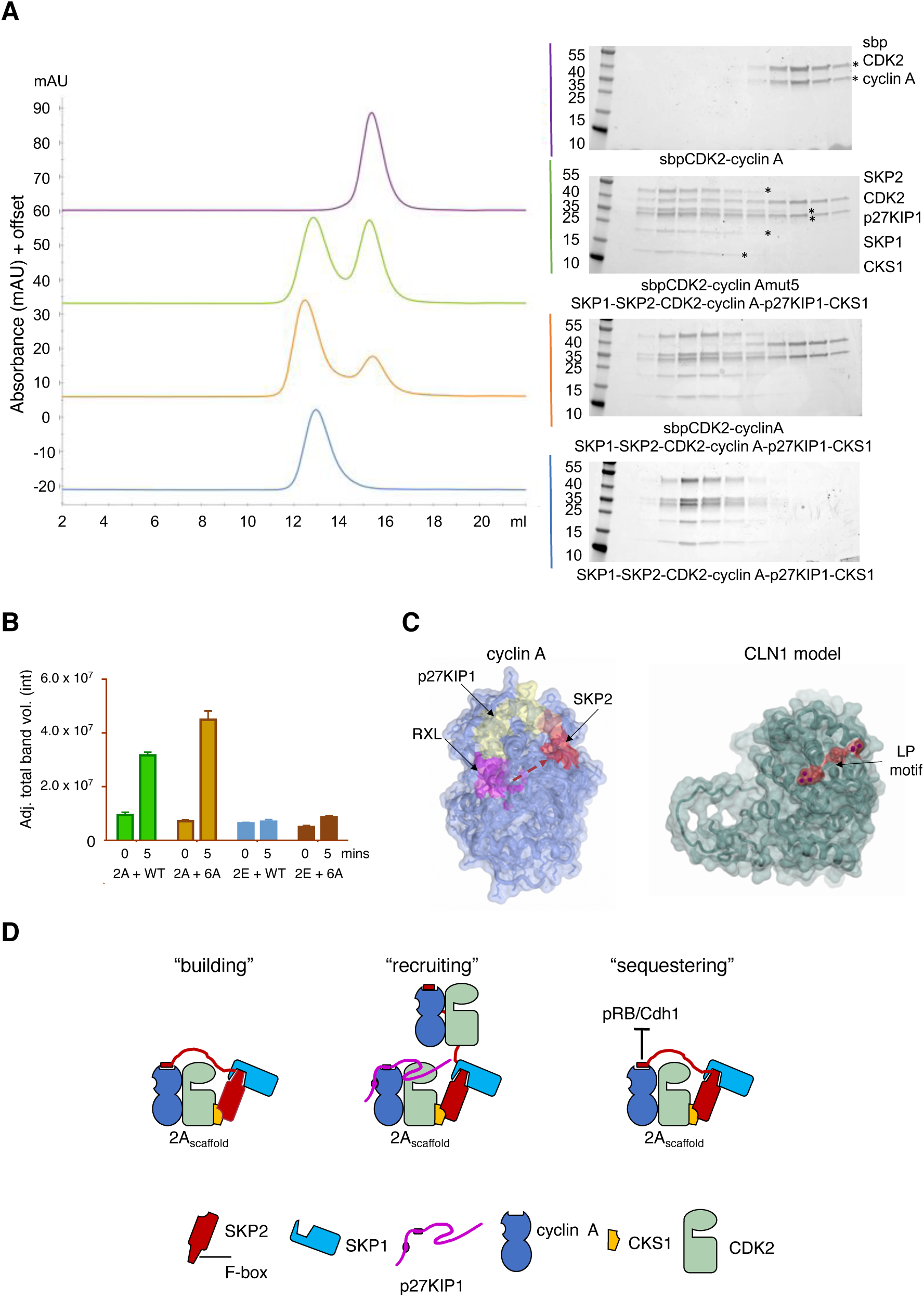
The cyclin A N-terminal cyclin box exploits an extended surface to interact with p27KIP1 and SKP2. (**A**) **A**) p27KIP1 association with CDK2-cyclin A can drive the formation of an octameric (CDK2-cyclin A)_2_-SKP1-SKP2-CKS1-p27KIP1 complex. p27KIP1 was first phosphorylated with an excess of CDK2-cyclin A and then purified by SEC before being incubated with equimolar SKP1-SKP2 and an excess of CKS1. This hexameric complex was then further purified by SEC before being incubated with either SBP-tagged CDK2-cyclinA or SBP-CDK2-cyclinAmut5. (**B**) The SKP2 CAIM recruits CDK-cyclin A to promote p27KIP1 phosphorylation. An intact cyclin A-SKP2 interaction recruits a catalytic CDK-cyclin module to phosphorylate p27KIP1 within a SKP1-SKP2-CDK2-cyclin A-CKS1-p27KIP1 complex. WT substrate, CDK2^D127A^-cyclin A-p27KIP1-SKP1-SKP2-CKS1; 6A substrate, CDK2^D127A^-cyclin A-p27KIP1-SKP1-SKP2_6A-CKS1. (**C**) p27KIP1 and SKP2 binding to cyclin A. p27KIP1 binding surface (yellow) identified from PDB 1JSU; SSMS site, red. Dotted red arrow, the proposed SKP2 binding path. *S. cerevisiae* CLN2 shares the docking site that recognizes the RXL motif (the “HP region” (Bhaduri et al., 2015)) with cyclin A and an adjacent surface (red), that recognizes LP-motif containing substrates. The CLN2 structure was generated by implementing MODELLER (Webb and Sali, 2017) in HHpred (Zimmermann et al., 2018) and then superimposed on the structure of cyclin A. (**D**) Proposed roles for the cyclin A-SKP2 interaction in regulating the cell cycle.

To test whether the recruitment of a catalytic CDK2-cyclin A via the CAIM-cyclin A interaction impacts the kinetics of phosphorylation of p27KIP1, we next compared the rate of phosphorylation of p27KIP1 bound within complexes that incorporate catalytically inactive (CDK2^D127N^)-cyclin A together with either CKS1-SKP1-SKP2 or CKS1-SKP1-SKP2_6A. These complexes were incubated under initial rate conditions with either CDK2-cyclin A or CDK2-cyclin E (Figure 4B). As expected, CDK2-cyclin E did not distinguish the two complexes, and p27KIP1 within each complex was phosphorylated at an equivalent initial rate. However, the initial rate of p27KIP1 phosphorylation was different when p27KIP1 was presented in the authentic complex than in the context of the SKP2-6A mutant. This result supports a model in which recruitment of CDK2-cyclin A to the hexameric substrate results in a relatively long-lived complex. Such a complex would be primed for rapid initial phosphorylation, with the slower turnover in steady state kinetics observed here being explained by a relatively slow product release step.

## Discussion

Cell cycle transition from G1 to S phase is tightly regulated by the sequential activation of CDK2 by cyclin E and then by cyclin A. p27KIP1 is both a CDK2 inhibitor and a CDK2 substrate and the balance of these activities ensures appropriate cell cycle progression. Signals to arrest the cell cycle during G1 are executed through p27KIP1-mediated CDK2 inhibition but, following checkpoint resolution, p27KIP1 is phosphorylated on T187 by CDK2-cyclin E and CDK2-cyclin A targeting it for ubiquitination by the SCF^SKP2^ E3 ubiquitin ligase and subsequent degradation by the proteasome. SKP2 levels are low in early G1 and increase late in G1 (Bassermann et al., 2014).

To characterize the molecular interactions that distinguish these various CDK2-containing complexes we set out to identify the SKP2 binding site on cyclin A. We show that the SKP2 binding site on cyclin A (the “SSMS site”), distinguishes cyclin A from cyclin E and is distinct but overlapping with the cyclin A binding site for p27KIP1. We also note that the *S. cerevisiae* LP docking site is co-located on this surface of the cyclin N-CBF (Bhaduri et al., 2015). Indeed, a homology model of Cln2 indicates that Cln2 R109 (part of the R109, L112 and R113 LP docking site), is equivalent to M246 of the SSMS motif (Figure 4C). Our study also revealed that sequences that do not conform to the canonical RXL motif can bind to the cyclin recruitment site. This finding suggests that the recruitment site might be important for the cyclin localization of a larger number of proteins than might have been inferred by sequence analysis. Taken together, these results reveal that the N-terminal cyclin box fold has an extended role in binding regulators. They also suggest that the cyclin A sequence and structure are finely tuned to optimize interactions with multiple binding partners. This site of direct interaction between SKP2 and cyclin A, in addition to the interaction between CKS1 and SKP2 shared with other CDK-cyclin modules, strengthens the cyclin A-SKP2 association and might explain why SKP2 was originally characterized as a component of a pentameric complex containing cyclin A and not cyclins B or E (Figure 4D “building”), (Zhang et al., 1995).

Furthermore, we show that the cyclin A SKP2 binding site permits the recruitment of CDK-cyclin A but not CDK-cyclin B or cyclin E complexes to perform a catalytic function and promote the phosphorylation of p27KIP1 on T187, a modification that is a pre-requisite for its subsequent degradation by the SCF^SKP2^ complex (Figure 4D “recruiting”). This activity is unique to cyclin A, whereas p27KIP1 recruitment to SKP2 is dependent on CKS1 and independent of the identity of the CDK-cyclin to which it is bound. Thus, our results agree with a previous study in which it was shown that the ability of SKP2 to bind to cyclin A could be separated functionally from its ability to mediate p27KIP1 degradation (Ji et al., 2006). Such a model might also explain earlier studies in G1-enriched cell cultures, where it was found that CDK2-cyclin A overexpression was more efficient at promoting p27KIP1 ubiquitination than increasing the expression of CDK2-cyclin E and SKP2 (Zhu et al., 2004).

However, it should also be noted that the SKP2 N-terminal sequence binds to other cell cycle regulators. Its occlusion within a pentameric SKP1-SKP2-CDK2-cyclin A-CKS1 complex may sequester it away from alternative interactions that might execute some of its functions or regulate its activity and thus perform an additional regulatory role. For example, cyclin A but not cyclin E competes with pRB and Cdh1 for binding to the SKP2 N-terminal sequence (Bashir et al., 2004; Wei et al., 2004). This interaction between pRB and SKP2 has been proposed as a key mechanism mediating G1 arrest (Ji et al., 2004). Expression of cyclin E in early G1 will not antagonize this interaction, but as the cell moves through to late G1, cyclin A levels rise and SKP2 can now be assembled into an alternative complex with SKP1, CDK2, cyclin A and CKS1. This complex will both hinder the ability of pRB to stabilize p27KIP1 and prime SKP2 for its alternative role to mediate p27KIP1 degradation. We hypothesize that both events could relieve cell cycle control from pRB dependency and promote S-phase entry. The SKP2 N-terminal sequence is also phosphorylated (at S64, S72 and S75) and acetylated (at K68 and K71), and it might be hypothesized that anchoring the SKP2 sequence on the surface of cyclin A affects the accessibility of these residues to post translation modification, thereby modulating its activity (Figure 4D, “sequestering”). Perturbations to these regulatory pathways may contribute in part to cancer progression seen in cells that overexpress cyclin A (Zhang et al., 1995), as the interactions that mediate these regulatory pathways would be in direct competition with unphosphorylated p27KIP1 for binding to CDK2-cyclin A as has been previously proposed (Ji et al., 2006; Lu et al., 2014).

The cyclin recruitment site that binds substrates and inhibitors containing an RXL motif is well characterized and illustrates the principle that a conserved site that is able to dock short peptide motifs with varying affinities offers a hub to integrate various signaling pathways. A similar principle orchestrates CDK9-cyclin T regulation by various activators and inhibitors that bind to overlapping surfaces on the C-terminal CBF (Gu et al., 2014; Schulze-Gahmen et al., 2013). We demonstrate that the SKP2 and p27KIP1 cyclin A binding sites overlap permitting integrated regulation of activity. To what extent this principle will apply to any additional cyclin A protein binding sites and to what extent the SKP2 binding site or these other sites will further distinguish cyclins A, B, D and E awaits further identification and characterization of larger CDK-cyclin containing complexes.

## Methods

### Protein expression and purification

Full-length human GSTCDK2 co-expressed with *S. cerevisiae* GSTCAK1 from a pGEX-6P-1 vector (Brown et al., 1999), (Brown et al., 2015)(Brown et al., 2015)and untagged human cyclin A2 (residues 174-432) (Brown et al., 1999), hexahistidine-tagged bovine cyclin A (residues 174-432) (Brown et al., 1995) and full-length human CKS1 (Brown et al., 2015) all cloned into the pET21d vector; and full-length p27KIP1 (residues 1-198, wild-type and T187A mutant), p27KIP1M (residues 1-106), p27KIP1S (residues 23-106) (Hallett et al., 2017), and p27KIP1MS (residues 23-51, this paper) as GST-fusions from a modified pGEX-6P1 vector, were all expressed in *E. coli* BL21 (DE3) pLysS cells and purified as previously described. Briefly, GST-p27KIP1 constructs, bovine cyclin A_His6_ and human CKS1_His6_ were all purified by an initial affinity chromatography step exploiting their respective tags. The GST constructs were subsequently digested with 3C protease to cleave the tag. All constructs were subject to a second SEC step. N-terminal Avi-tagged CKS1 (residues 5-73) was expressed as an N-terminally His_6_-tagged protein from the pET21d vector. It was purified by affinity chromatography, subsequent 3C protease cleavage to remove the His_6_ tag, then re-application on to the affinity column to remove any uncleaved protein before a final SEC step. The global human cyclin A mutant sequence was synthesized by IDT, sub-cloned into an IDTsmart vector and then PCR amplified and inserted into pET21d to generate an untagged protein. Other human cyclin A mutants were prepared either starting from this template or the authentic sequence by site-directed mutagenesis (SDM). All CDK2-cyclin A complexes were purified following the method described in Brown et al., 1999 (Brown et al., 2015; Brown et al., 1999)(Brown et al., 2015; Brown et al., 1999). Briefly, GST-CDK2 was first loaded onto a glutathione affinity column and washed to baseline. This modified column was then used to affinity purify untagged cyclin A. The glutathione eluate (eluted with 20 mM HEPES, NaCl 300 mM, DTT 1mM, 20 mM glutathione, pH7.6) was incubated with 3C protease to release the CDK2–cyclin A from GST, which was subsequently further purified by SEC and glutathione affinity chromatography to remove any co-eluting GST. All steps from re-suspension of the bacterial cell pellets were carried out in buffer containing 20 mM HEPES pH 7.6, 300 mM NaCl, 1mM DTT. The CDK2 D127A mutant was introduced into a modified pET3d vector that co-expresses GST-CDK2, His-tagged bovine cyclin A and untagged p27KIP1 and the complex subsequently purified by sequential glutathione affinity and size exclusion chromatography steps. Human cyclin E1 (residues 96-378) was cloned into a modified pACEBac1 vector containing an N-terminal MBP-3C tag. PCR of cyclin E1 (96-378) from a modified pGEX vector was conducted using primers: 5’-TTCCAGGGGCCCATGGGCATTATTGCACCATCCAGAGGCTCCC-3’ (Forward) and 5’-GTTAGCAGCCACTAGTTCAGGCTTTCTTTGCTCGGGCTTTGTC-3’ (reverse). pACEBac1_MBP-3C-cyclin E was then transformed into DH10_YFPEMBac *E. coli* for virus transposition, with appropriate clones selected through blue/white screening. 20 mL overnight cultures (in Insect Xpress media (Lonza)) of selected clones were grown. The virus was then extracted by isopropanol precipitation and amplified twice in Sf9 cells. For cyclin E expression, the virus was transfected into Sf9 cells at approximately 1.5 million cells/mL and grown for 72 hours at 28 °C with gentle mixing. Following incubation cells were harvested via centrifugation (2600 xg, 15 mins, 4 °C) and resuspended in 20 mM HEPES pH 7.6, 300 mM NaCl, 1mM DTT, supplemented with protease inhibitors. Cells were lysed by sonication and clarified by centrifugation (100000 xg). The soluble fraction was incubated with GST-CDK2 and then purified by affinity chromatography using Sepharose 4B GST resin beads (GE Healthcare) and subsequently eluted in 20 mM HEPES, 300 mM NaCl, 1 mM DTT, 20 mM glutathione, pH7.6. Following 3C protease treatment CDK2-cyclin E was buffer exchanged using SEC (Superdex 200 10/300) and then loaded onto a GST-trap column (5ml, GE Healthcare) and then onto an amylose resin (New England Biolabs). A final SEC step (Superdex 200 10/300) was performed in order to remove any GST and MBP contamination. Full-length SKP2 was co-expressed as a GST fusion (cleavable with 3C protease) with untagged full-length SKP1 from a modified pET3d backbone. Each protein was expressed from separate transcripts. A GSTSKP2N-SKP1 complex was expressed from a pGEX-6P-1 vector in *E. coli* BL21 (DE3) pLysS cells (Agilent Technologies). This construct encodes SKP2 residues 1-140 (SKP2N) and SKP1 (residues 1-163, with internal deletion of residues 38-43 and 69-83 as described by Schulman *et al*, (Schulman et al., 2000)). The two proteins are translated from a single transcript, the SKP1 initiating at an internal ribosome binding site. This construct was used as the template to make the SKP1-SKP2N_4A (L32A, L33A, S39A and L41A), and SKP1-SKP2N_6A (SKP2_4A and W22A and W24A) mutants by site-directed mutagenesis (SDM) and the construct starting at F20 (SKP1-SKP2N_Δ20) by PCR. N-terminal SKP2 peptides were either synthesized (FTWGWDSSKTSELLSGMGVSALE) and modified by N-terminal acetylation and C-terminal amidation (Severn Biotech Ltd) or generated by SDM (SKP2 residues 20-51 and 20-64) starting with the GSTSKP2N-SKP1 expression construct. For the HTRF assay presented in Supplementary Figure S7, the SKP2 C-terminal sequence was mutated by SDM to replace F393 with glycine and serine (SKP2 F393GS) and co-expressed and purified with SKP1. The construct using the pGEX-6P-1 backbone is based on that described in Schulman *et al*, (Schulman et al., 2000) that encodes SKP2 residues 101-436 and SKP1 (residues 1-163, with internal deletion of residues 38-43 and 69-83). The SKP2 phosphorylation site mutant (S64A) was prepared by SDM using the GSTSKP2-SKP1 co-expression construct in the modified pET3d backbone described above that encodes full-length SKP2. E2 R1 and all the SKP1-SKP2 expression constructs described above were purified by sequential glutathione affinity and SEC chromatography steps in 20 mM HEPES, 300 mM NaCl, 1 mM DTT. The p107 substrate peptide was prepared as described in (Brown et al., 2015). Briefly, the protein was expressed as a GST-fusion from the pGEX6P-1 backbone in *E. coli* Rosetta (DE3) pLysS cells at 20 °C overnight subsequent to induction with 0.1 mM IPTG. It was first purified by a glutathione-affinity step and then subsequent to tag removal following incubation with 3C protease further purified on a MonoS ion-exchange (GE Healthcare) column equilibrated into 25 mM MES pH 6.5. The peptide was stepped into 25 mM MES, 0.5M NaCl, pH6.5 for peptide rapid release. Fractions containing the peptide were confirmed by SDS–PAGE, pooled, concentrated to a final peptide concentration between 1 and 2 mg ml-1 and then flash frozen and stored at ^-^80 °C in 100 μl aliquots. Protein concentrations were measured by A_280_ nm on a Nanodrop2000 (Thermo Fisher Scientific) using calculated extinction coefficients (http://web.expasy.org/protparam/).

### Isothermal titration calorimetry (ITC)

ITC was performed using a Malvern Microcal PEAQ-ITC. Proteins were equilibrated in HEPES 20 mM pH 7.97, NaCl 300 mM, TCEP 0.5 mM. The concentrations of CDK2-cyclin A or cyclin A in the cell and SKP1-SKP2N in the syringe for each experiment are provided in the accompanying Table Legends. Protein concentrations (for this and other biophysical techniques) were measured by A_280_ nm on a Nanodrop2000 (Thermoscientific) using calculated extinction coefficients (http://web.expasy.org/protparam/). Experiments (including ligand dilution controls) were carried out at 25°C. The titrations consisted of 20 injections, an initial injection of 0.5 μL and then 19 injections of 19.5 μL with 180 seconds between injections. The stirring speed in the reaction cell was 1000 rpm. Data were analyzed using non-linear least squares regression using MicroCal PEAQ analysis software. Drifts in the baseline were corrected during data analysis. As detailed in Table 1, for characterization of authentic interactions and mutants that were characterized further by other orthogonal techniques, K_d_ values are presented as the average of two independent biological replicates and SD of the population. For other mutants, K_d_ values were derived from one measurement where the error represents the fitting error directly derived from the thermogram. Significance was calculated at p < 0.05 using a Student’s t-test.

### Analytical size-exclusion chromatography

Size exclusion chromatography was performed using a Superdex 200 (10/300, GE Healthcare) on an AKTA Pure at controlled temperature of 4 °C, pre-equilibrated in HEPES 20 mM pH 7.5, NaCl 300 mM, DTT 1 mM. Each complex was reconstituted *in vitro* in 0.4 ml of HEPES 20 mM pH7.5, NaCl 300 mM, DTT 1 mM and glycerol 10%, and incubated for 1 hour at 4⁰C prior to elution. The complexes were combined at a molar ratio of 1:1.5:2.25 corresponding to SKP1-SKP2 (4 µM):CDK2-cyclin A (6 µM):CKS1 (9 µM). For reconstitution of the p27KIP1-containing complex the ratio was adjusted to 1:1.5:2.25:3.375 using SKP1-SKP2 (4 µM):CDK2-cyclinA (6 µM):p27KIP1S (9 µM):CKS1 (13.5 µM). Calibration of the Superdex 200 (10/300) was performed using β-Amylase (200 kDa), albumin (66 kDa) and carbonic anhydrase (29 kDa) protein molecular weights standards (Sigma Aldrich). Corresponding fractions for each chromatogram between 12 and 18 ml with an interval of 0.5ml were analyzed by SDS-PAGE (Biorad) and Instant blue coomassie staining (Expedeon).

### Surface plasmon resonance (SPR)

SPR analysis was carried out using a Biacore S200 (GE Healthcare). The sensor chip CM5 surface was functionalized using an anti GST tag antibody by amine-coupling. Routinely, ligand capture on flow cell number 2 was carried out for 60 s followed by multiple buffer injection to remove the excess of inactive protein, and the intensity was normalized against flow cell number 1 (GST control as ligand to correct for non-specific interactions and bulk effects). Experiments were performed in HEPES 20 mM pH7.4, NaCl 150 mM Tween20 0.01% at 20 °C. GSTp27KIP1M was captured on the functionalized surface of flow cell 2 for 60 seconds at a flowrate of 5 µl min^-1^. CDK2-cyclin A wild-type and mutants were then titrated as analytes using a 4-fold dilution series and then the chip was regenerated using glycine pH 2.1 for 60 seconds. The sample contact time (60 s) was measured using high performance settings followed by 240 seconds of dissociation time at 30 µl/min. Two independent (biological replicate) experiments were performed. Sensorgrams were analyzed using Biacore S200 Evaluation Software 1.0 (GE Healthcare) and the curves were fitted using 1:1 binding kinetics.

### Homogenous time-resolved fluorescence assay (HTRF)

In this format the HTRF assay measures the interaction between GSTSKP2-SKP1 and N-terminal Avi-tagged CKS1. Optimized buffer and detection reagent conditions were 3 nM Tb-anti-GST antibody, 50 nM SA-XL665, in 50 mM HEPES pH 7.5, 100 mM NaCl, 1 mM DTT, 0.1 mg/ml BSA, 1% DMSO. Experiments were carried out at GSTSKP2-SKP1 or GSTSKP2_C2-SKP1 concentrations of 1, 5 or 25 nM into which N-AviCKS1 was titrated from 1 to 1000 nM. Reactions were set up in a volume of 20 µl in 384 well plates and incubated for 30 mins at 4 °C. Samples were excited using a wavelength of 337 nm and emission spectra measured at 620 nm and 665 nm using a PHERAstar plate reader (BMG Labtech). Binding curves were plotted using GraphPad Prism 6 from which the K_d_s were determined.

### Differential scanning fluorimetry

A protein/Sypro orange mix containing 5 μM protein and a 1:5,000 dilution of dye in DMSO (as supplied by Sigma Aldrich) was prepared just before plate setup in 20 mM HEPES pH 7.5, 150 mM NaCl and 0.5 mM TCEP. 14.5 μL of the protein/Sypro orange mix was aliquoted into 384-well plates and sealed. Each experiment was done in triplicate. Thermal melting experiments were carried out using a QuantStudio Real Time PCR machine (Applied Biosystems). The plates were first equilibrated at 25°C for 2 min in the PCR machine before starting the thermal melting experiment upon which the plates were heated at 0.05°C per second from 25 to 99°C. The fluorescence intensity was recorded using the ROX filters under continuous collection. Raw fluorescence data were extracted from the QuantStudio Real-Time PCR software and analysed using the Applied Biosystems Protein Thermal Shift software. Calculated derivative T_m_ values for each of the three repeats were extracted and the average values calculated.

### Hydrogen-deuterium exchange mass spectrometry

HDX-MS experiments were carried out using an automated HDX robot (LEAP Technologies, Fort Lauderdale, FL, USA) coupled to an M-Class Acquity LC and HDX manager (Waters Ltd., Wilmslow, Manchester, UK). Analysis of the CDK2-cyclin A-p27KIP1 and CDK2-cyclin A-SKP1-Δ20SKP2 complexes was carried out separately and so were plotted individually. All samples were diluted to 10 µM in equilibration buffer (50 mM potassium phosphate buffer pH 7.4) prior to analysis. 30 µL sample was added to 135 µl deuterated buffer (10 mM potassium phosphate buffer pH 7.4) and incubated at 4 °C for 0.5, 1, 2, 10 or 60 min. Following the labelling reaction, samples were quenched by adding 50 µl of the labelled solution to 100 µl quench buffer (50 mM potassium phosphate,0.05% DDM pH 2.2) giving a final quench pH ∼2.5. 50 µl of quenched sample (ca. 24 pmol) were passed through an immobilized Ethylene Bridged Hybrid (BEH) pepsin column (Waters Ltd., Wilmslow, Manchester, UK) at 500 µl min-1 (20 °C) and a VanGuard Pre-column Acquity UPLC BEH C18 (1.7 µm, 2.1 mm X 5 mm, Waters Ltd., Wilmslow, Manchester, UK) for 3 mins in 0.3% formic acid in water. The resulting peptic peptides were transferred to a C18 column (75 µm X 150 mm, Waters Ltd., Wilmslow, Manchester, UK) and separated by gradient elution of 0-40 % MeCN (0.1 % v/v formic acid) in H_2_O (0.3 % v/v formic acid) over 7 mins at 40 µl min-1. Trapping and gradient elution of peptides was performed at 0 °C. The HDX system was interfaced to a Synapt G2Si mass spectrometer (Waters Ltd., Wilmslow, Manchester, UK). HDMSE and dynamic range extension modes (Data Independent Analysis (DIA) coupled with IMS separation) were used to separate peptides prior to CID fragmentation in the transfer cell (Cryar et al., 2017). HDX data were analyzed using PLGS (v3.0.2) and DynamX (v3.0.0) software supplied with the mass spectrometer. Restrictions for identified peptides in DynamX were as follows: minimum intensity: 1000, minimum products per MS/MS spectrum: 5, minimum products per amino acid: 0.3, maximum sequence length: 25, maximum ppm error: 5, file threshold: 3/3. Following manual curation of the data, PAVED was used (Cornwell et al., 2018) to determine areas of the protein showing significant differences in exchange between the states. Briefly, for each time point and each amino acid residue in the sequence, the mean relative fractional uptake was calculated from all peptides covering that amino acid where the amino acid is not at the N-terminus of the peptide. Analysis of variance and post-hoc Tukey tests were used to determine significant differences (p<0.05). For each region of significant difference, a representative peptide was chosen for the following discussion.

### Small angle X-ray scattering and model building

The SKP1(Δ38-43, Δ69-83)-SKP2(20-140-Δ60-74)-CDK2-cyclin A complex was purified by size exclusion chromatography in 20 mM HEPES 150 mM NaCl 1 mM DTT pH 7.0 at a final concentration of 5mg/ml. On the BL21 beamline at the Diamond Light Source the complex was injected though a HPLC-size exclusion column collecting 490 individual SAXS frames (Supplementary Table 3). Fractions were analyzed at 10°C, using an X-ray wavelength of 12.4 KeV at a detector distance of 4.014 m, resulting in the momentum transfer range of 0.004 Å^−1^ < q < 0.4420 Å^−1^. The average and the median were calculated from peak fractions subtracted from selected buffer frames of the column void. Data were scaled, merged and analyzed using Scatter3 (Rambo and Tainer, 2015). To generate the model, a SKP2 fragment (28-45) was built in MOE (2018.1), energy minimized and subject to conformer generation. A randomly selected conformer was docked to cyclin A (PDB 1VIN) in MOE using the MRAIL and SSMS motifs as targeted binding sites. Best solutions were subject to molecular dynamics for refinement and the resulting model with the best interface area (as analyzed by PDBePISA (v1.52)) was used as a starting model to extend the peptide N-terminally to the full fragment 17-45 (ATSFTWGWDSSKTSELLSGMGVSALEKEE). The extended model was refined in molecular dynamics simulation. A 35 dummy atom linker was introduced to connect the peptide to SKP1-SKP2(Fbox) (PDB 2AST). Ab initio molecular bead models were generated using DAMMIF (Franke and Svergun, 2009). Rigid body modelling carried out within CORAL(Petoukhov et al., 2012) gave a Chi square fitting of 1.45 with the experimental data. The volume-adjusted (‘filtered’) average of these models (damfilt.pdb) was converted into a map using molmap and visualized in ChimeraX(Goddard et al., 2018).

### Crystallization and structure determination

CDK2-cyclin Amut 5 was concentrated to 5.6 mg ml^-1^ in HEPES 20 mM pH 7.6, NaCl 150 mM, DTT 1 mM. Crystallization trials were set up at 4 °C using the sitting drop vapor diffusion method. Crystals were grown in condition B10 of the JCSG+ crystallization screen (Molecular Dimensions) composed of 200 mM K/Na Tartrate, 20 % PEG 3350. They were subsequently harvested using fibre loops and flash-cooled in liquid nitrogen, using 20 % PEG400 as cryo-protectant. X-ray diffraction data were collected at 100 K at the Diamond Light Source (Didcot, UK), beamline IO4-1 operating at a wavelength of 0.92 Å. The automatic molecular replacement program MOLREP (within the CCP4 suite (Potterton et al., 2018)) was used to calculate molecular replacement solutions and the correlation coefficients. Model building was done using Coot (Emsley et al., 2010). The final model has 0.3 %, 1.8 % and 1.4 % defined as Ramachandran, sidechain or RSRZ outliers (wwPDB X-ray structure validation report). Structural figures were prepared using the CCP4 molecular graphics program CCP4MG (McNicholas et al., 2011). Statistics generated from X-ray crystallography data processing, refinement, and structure validation are displayed in Supplementary Table S1.

### Kinase assays

Non-phosphorylated p27KIP1 substrate complexes were assembled by incubating co-expressed CDK2^D127A^-cyclin A-p27KIP1 with an excess of either SKP1-SKP2-CKS1 or SKP1-SKP2_6A-CKS1 and subsequently purified using a Superdex 200 column equilibrated in 20 mM Tris-HCl pH 7.8, 300 mM NaCl, 0.5 mM TCEP, 5% glycerol. Kinase assays were carried out in SEC buffer supplemented with 0.75 mM ATP, 2 mM MgCl_2_, 0,1mg/ml BSA. 2.5 nM CDK2-cyclin A was incubated with a 2170 nM of substrate complex at 15°C and the reaction was stopped after 5, 10 and 20 minutes by addition of Laemmli buffer at 95°C for 2 min, and subsequently analyzed by SDS-PAGE (Criterion, Biorad). CDK2-cyclin E was initially assayed using the same conditions and then subsequently assayed at 0 and 5 mins with its concentration adjusted to 5 nM. Protein samples were transferred and western blotted using anti phospho-T187 p27KIP1 antibody (Merck, 06-996). Band intensity was revealed using ECL reagent (BioRad) and quantified using background adjusted total band volume with ImageLab software (BioRad). Assays were carried out with three biological replicates of catalytically active CDK2-cyclin A and CDK2-cyclin E as control. Other kinase assays were carried out as described (Brown et al., 2015). Briefly, phosphorylated CDK2–cyclin A, selected CDK2-cyclin A mutants, CDK2-cyclin A-SKP1-SKP2 (full-length SKP1 and full-length wild-type SKP2 and S64A mutant) and CDK2-cyclin A-p27KIP1M (residues 1-106) were assayed using the ADP-Glo^TM^ assay (Promega) using a p107 peptide (peptide SPIK+RXL: IGSGVLFQGPLGSMHPRVKEVRTDSGSLRRDMQPLSPISVHER WSSATAGSAKRRLFGEDPPKEMLMDKW (Brown et al., 2015)) essentially as described by the manufacturers. In brief, in a final volume of 5 μl, reactions were carried out at room temperature in 40 mM Tris-HCl pH 7.5, containing 20 mM MgCl_2_, 0.1 mg ml ^-1^ BSA, and 1.5 nM CDK2–cyclin A or CDK2-cyclin A-SKP1-SKP2 complex. Peptide dilutions (94 to 0 μM) were added to the CDK–cyclin solution and the reactions were initiated by the addition of 75 or 25 μM ATP (as indicated in Supplementary Table S2). All activity assays were performed in duplicate or triplicate (as indicated in Supplementary Table S2) in white low volume 384-well plates using a PheraStar plate reader (BMG). K_m_ and V_max_ values were obtained by fitting the data to the equation v = V_max_/(1 + (K_m_/[S]) using PRISM (GraphPad).

### Cell-based studies

HEK293T (female) cells were propagated in DMEM supplemented with 10% fetal bovine serum (FBS) (Corning Life Sciences) and 1% penicillin/streptomycin/L-glutamine (Corning Life Sciences). Cyclin A, cyclin A_mut5, SKP2_6A, SKP2_C2, and SKP2_6A_C2 cDNAs were inserted into a modified pcDNA3.1 vector containing an N-terminal FLAG tag by subcloning. SKP2 cDNA was inserted into the pcDNA3.1 vector containing an N-terminal FLAG/Strep tag. HEK293T cells were transiently transfected using polyethylenimine. Twenty-four hours after transfection, HEK293T cells were collected and lysed with lysis buffer (50 mM Tris pH 8.0, 150 mM NaCl, 10% glycerol, 1 mM EDTA, 50 mM NaF, and 0.2% NP-40) supplemented with protease and phosphatase inhibitors. Lysates were then immunoprecipitated with anti-FLAG antibody conjugated to agarose. Elution of the immunoprecipitation from the anti-FLAG agarose resin was carried out with 3xFLAG peptide. Immunoblotting was performed as previously described (Marzio et al., 2019). Briefly, samples were resolved under denaturing and reducing conditions using 4-12% Bis-Tris gels (NuPAGE®) and transferred to a PDVF membrane (Immobilon-P, Millipore). Membranes were blocked with 5% nonfat dried milk, then incubated with primary antibodies overnight at 4°C. After washing the membranes, secondary antibodies coupled with horseradish peroxidase were applied (Amersham-GE). Immunoreactive bands were visualized by enhanced chemiluminescence reagent (Thermo Fisher Scientific) and signal was acquired using ImageQuant LAS 400 (GE). The following antibodies were used for immunoblotting: anti-CUL1 (Santa Cruz Biotechnology, sc-11384, 1:500), anti-FLAG (Sigma, F7425, 1:2000), SKP2 (Invitrogen, 32-3300, 1:1000). Antibodies against CKS1, SKP1, cyclin A, cyclin B, CDK1 and CDK2 are homemade from the Michele Pagano lab.

## Acknowledgements

X-ray crystallography and SAXS was carried out with the support of Diamond Light Source on beamlines I04-1 (proposal MX13587) and BL21 (sm16970-2) respectively. We thank N. Schueller, M. Hoellerer and H. Ruddick (University of Oxford) for preparation and initial construct characterization and for assistance with preparation of the mutants; K. Cole, M. Martin, C. Jennings and A. Wittner for assistance with the SEC, ITC, mass spectrometry and cell-based studies respectively. H. Waller and O. Davies provided invaluable assistance to carry out the ITC and SEC-MALLS. B. Hao (U. Connecticut) for providing the SKP1-SKP2 co-expression plasmid. This research was supported by the Wellcome Trust (Grant Reference 063551); MRC (Grant References G0901526 and MR/N009738/1), Cancer Research UK (Grant Reference C2115/A21421), Newcastle Cancer Centre and the JGW Patterson Foundation and the BBSRC (Grant Reference BB/M012573/1). The LEAP sample handling robot used in the HDX-MS work was a kind donation from Waters UK. M.P. is an Investigator with the Howard Hughes Medical Institute and funded by grants from the National Institutes of Health to M.P. (GM136250 and CA76584). Research by S.A. reported in this publication was supported by the King Abdullah University of Science and Technology (KAUST).

## Author contributions

M.S. Construct design, protein production, biochemical assays, biophysical methods (SEC, SPR, ITC, SAXS) and X-ray crystallography.

B.C.M. Cloning, construct design, protein production, ITC and immunoprecipitation experiments.

D.W. Protein production

R.H., Construct design

L.W. and S.K. ADPGlo kinase assays

J.R. Selected HTRF assays

A.B. Data collection and structure determination

M.W.P. HDX-MS sample preparation

N.J.T. MD simulations and molecular docking

J.R.A. HDX-MS data collection and analysis

F.S. HDX data analysis

S.A. SAXS data analysis and interpretation

M.L. Conceptualization and investigation

M.P. Conceptualization, resources, supervision, and funding acquisition

M.S., J.A.E. and M.E.M.N. designed and supervised the project. All authors analyzed the data and B.M., M.S., M.E.M.N. and J.A.E. wrote the paper.

## Additional information

M.P. has financial interests in CullGen Inc. and Kymera Therapeutics. M.P. is on the SAB of CullGen Inc. and Kymera Therapeutics. M.P. is a consultant for CullGen Inc., Kymera Therapeutics, Exo Therapeutics, and SEED Therapeutics. J.R. is a current employee of Astex Pharmaceuticals. The other authors declare no competing interests. The authors declare no competing financial interest. Some work in the authors’ laboratory is supported by a research grant from Astex Pharmaceuticals. The coordinates and the structure factors have been deposited in the Protein Data Bank under ID code 6SG4. Other data are available from the corresponding author upon reasonable request. Abbreviations used: CAIM, cyclin A interacting motif; CBF, cyclin-box fold; CDK, cyclin-dependent kinase; CKI, cyclin-dependent kinase inhibitor; CKS1, cyclin-dependent kinases regulatory subunit 1; DSF, differential scanning fluorimetry; GST, Glutathione-S-transferase; HDX-MS, hydrogen-deuterium exchange mass spectrometry; ITC, isothermal titration calorimetry; LRR, leucine rich repeats; RMSD, root mean square deviation; SAXS, small angle X-ray scattering; SEC, size exclusion chromatography; SKP, S-phase kinase associated protein; SPR, surface plasmon resonance

---

**Supplementary Figure S1.**
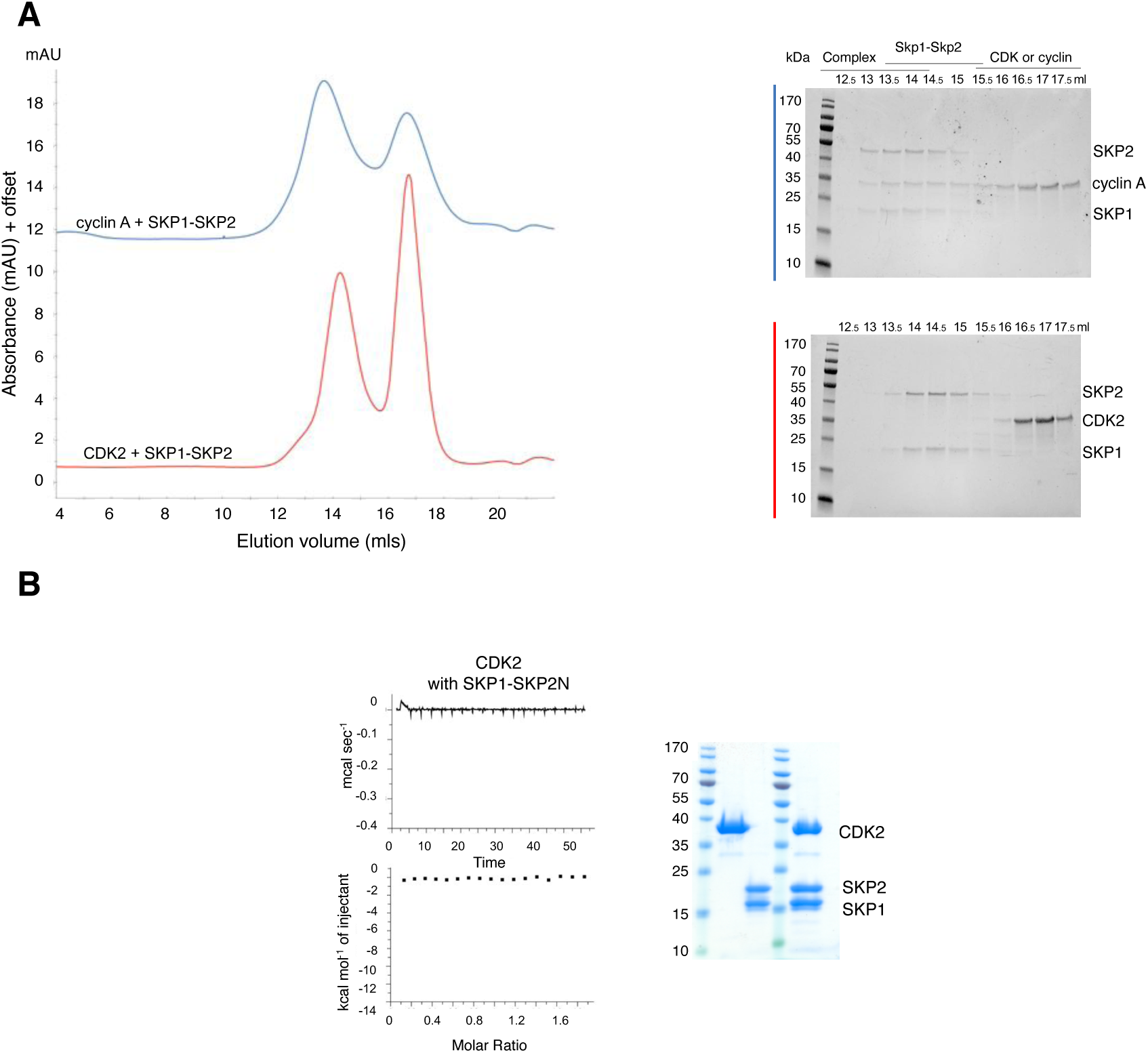
Characterization of the binding between SKP1-SKP2 and CDK2 and cyclin A. Related to Figure 1. (**A**) SKP1-SKP2 binds to cyclin A but not CDK2. SKP1-SKP2 was mixed with an excess of cyclin A (blue trace) or CDK2 (red trace) and analyzed by analytical size-exclusion chromatography. For each run comparable 0.5 ml fractions were analyzed by SDS-PAGE and visualized by InstantBlue staining. Chromatograms are to the same scale but have been offset on the y-axis to aid comparison. Cyclin A construct is bovine cyclin A residues 174-432. (**B**) SKP1-SKP2N does not bind to CDK2. Isothermal titration calorimetry (ITC) to measure the affinity of SKP1-SKP2N for monomeric CDK2. SDS-PAGE analysis confirms that post titration of SKP1-SKP2N and CDK2 all three proteins are intact (compare lanes 2 and 3 with lane 5). ITC reaction conditions are provided in Table 1. Chromatogram is representative of two experiments carried out using independently prepared proteins. The titration of monomeric CDK2 against SKP1-SKP2N was carried out once.

**Supplementary Figure S2.**
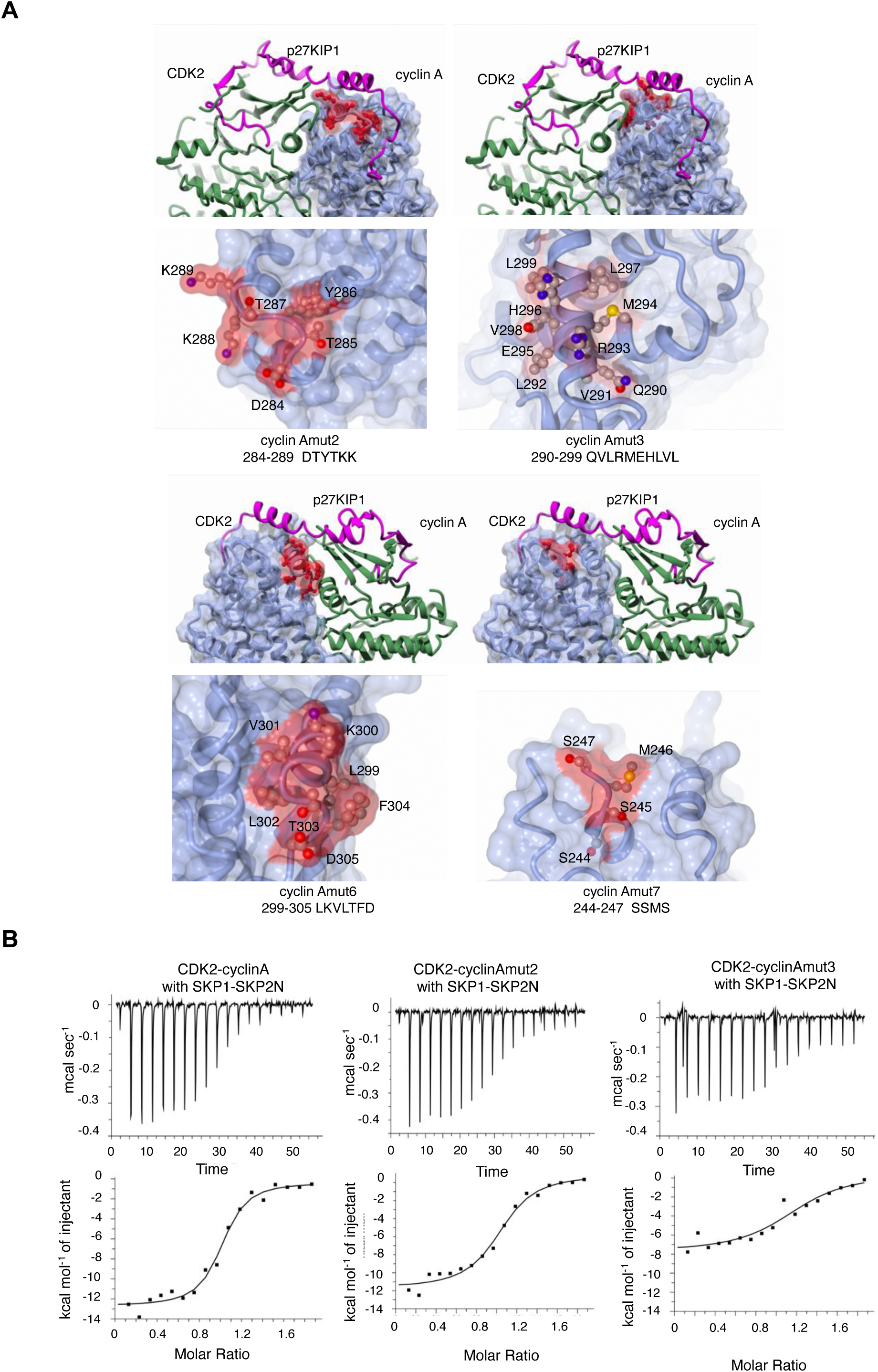
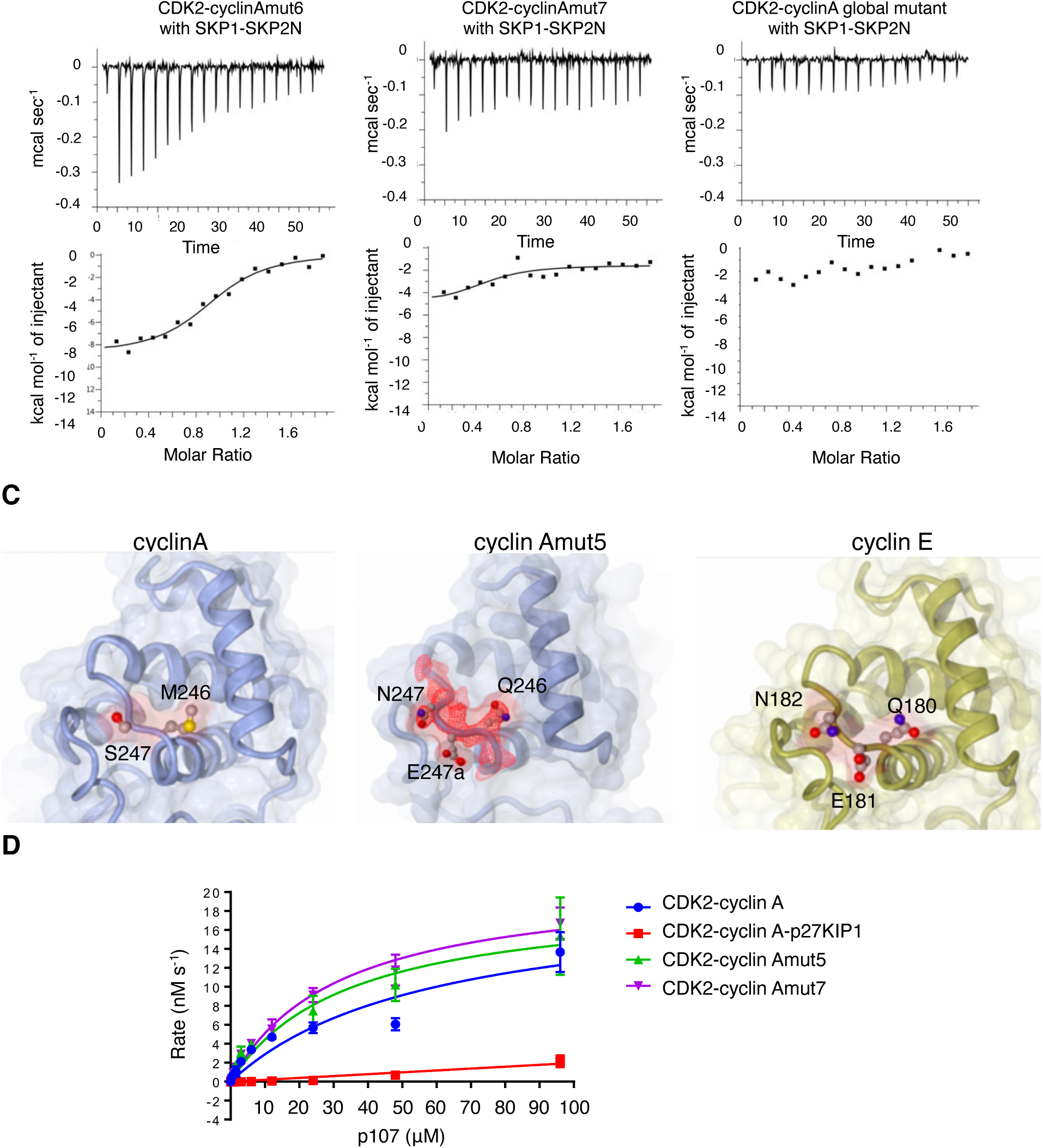
Characterization of cyclin A mutants. Related to Table 1. (**A**) Cyclin A mutant locations on the structure of a CDK2-cyclin A-p27KIP1 complex (PDB entry 1JSU). The cyclin A surface is rendered in blue, and the CDK2 and p27KIP1 folds in green and magenta respectively. Residues mutated in each of the cyclin A mutants are highlighted in red. (**B**) Isothermal titration calorimetry to measure the interaction between SKP1-SKP2N and mutant CDK2-cyclin A complexes. Thermodynamic parameters and binding constants are tabulated in Table 1. (**C**) Structure of cyclin Amut5. The structure of cyclin Amut5 is shown in the middle panel and compared to that of cyclin A (LHS panel, extracted from PDB entry 1QMZ, (Brown et al., 1999)) and cyclin E (RHS panel, extracted from PDB entry 1W98 (Honda et al., 2005)). Cyclin A residues M246 and S247 and cyclin E residues Q180, E181 and N182 (Uniprot entry P24864, numbered as Q165, E166 and N167 in PDB 1W98) are drawn in ball and stick mode. The electron density map that supports the cyclin Amut5 structure at the mutation site is drawn as a red mesh and contoured at 1σ. The location of the SSMS site on the cyclin A structure is also highlighted in red in Figure 1C. This structure adopts an unusual packing in the crystal in which CDK2 N-lobes undergo partial unfolding and domain exchange, but in which the structure of cyclin A is unperturbed. The structures of cyclin A, cyclin E, and cyclin Amut5 overlay closely in the region of the mut5 mutations, although cyclin E and cyclin Amut5 accommodate an insertion where cyclin E residues E181 and N182 replace cyclin A S247. This sequence insertion is one of only two insertions/deletions that distinguish their respective N-terminal cyclin box folds (CBFs). A superposition of cyclin A and cyclin Amut5 indicates that the insertion has not affected the overall cyclin A fold: the extra residue forms a bulge within a surface loop, returning to register at residue 249. As predicted from the cyclin E structure, residue E247a of cyclin Amut5 (analogous to cyclin E E181), orientates into solution and alters the charge of a cleft within cyclin A. (**D**) CDK2-cyclin A and CDK2-cyclin A mutants have comparable catalytic activity. CDK2-cyclin A activity was measured using the ADP-Glo^TM^ assay format towards a peptide substrate derived from the sequence of p107 that contains the sequence SPIK around the site of phosphor-transfer and an RXL recruitment site motif (sequence KRRL) (Brown et al., 2015). CDK2-cyclin A-p27KIP1(p27KIP1 residues 1-106) is not active in this assay (red trace). Kinetic parameters were derived using PRISM (GraphPad) and are presented in Supplementary Table S2. Protein kinase assays were performed in triplicate and error bars correspond to the range of values.

**Supplementary Figure S3.**
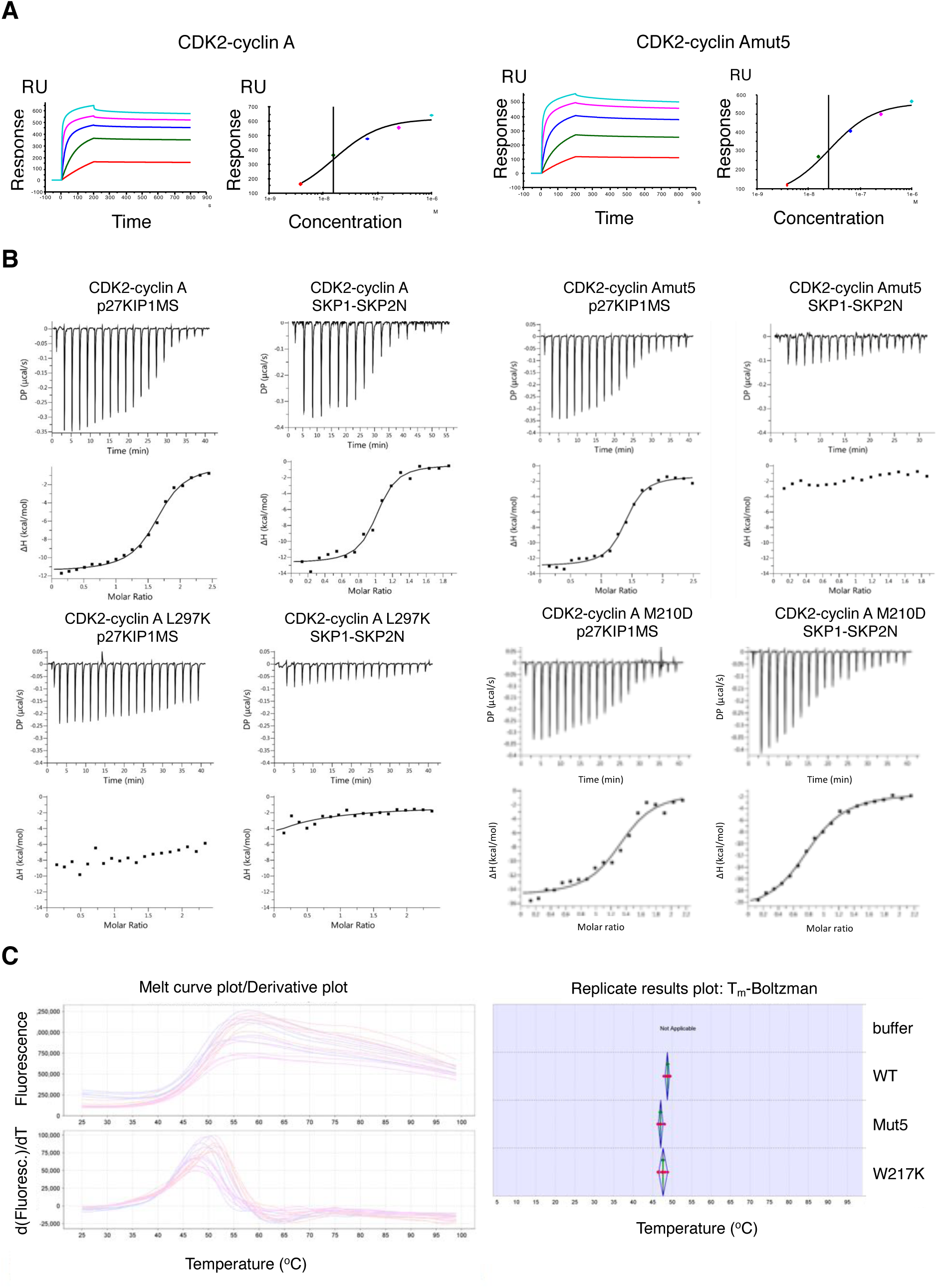
Cyclin A mutations distinguish the SKP2 and p27KIP1 binding sites. Related to Figure 2. (**A**) Surface plasmon resonance to measure the interaction between p27KIP1 and CDK2-cyclin A and CDK2-cyclin Amut5. GST-p27KIP1M (residues 1-106) was bound to a CM5 sensor-chip functionalized with anti-GST antibody and the CDK2-cyclin A complexes were flowed over at increasing concentrations. K_d_ values were derived using Biacore S200 evaluation software. (**B**) Isothermal titration calorimetry (ITC) to measure the interaction between SKP1-SKP2N or p27KIP1MS (residues 23-51) and mutant CDK2-cyclin A complexes. K_d_ values and reaction conditions are tabulated in Table 1. SPR sensograms and ITC thermograms are representative of two repeats using independently prepared protein samples. (**C**) Differential scanning fluorimetry to assess the stability of the cyclin A fold. Raw fluorescence data were extracted from the QuantStudio Real-Time PCR software (LHS panel) and analyzed using the Applied Biosystems Protein Thermal Shift software (RHS panel). Derivative T_m_ values for each complex were extracted and the average values from 3 technical repeats calculated. Red curves, CDK2-cyclin A, magenta curves, CDK2-cyclin Amut5, blue curves CDK2-cyclin AW217K.

**Supplementary Figure S4.**
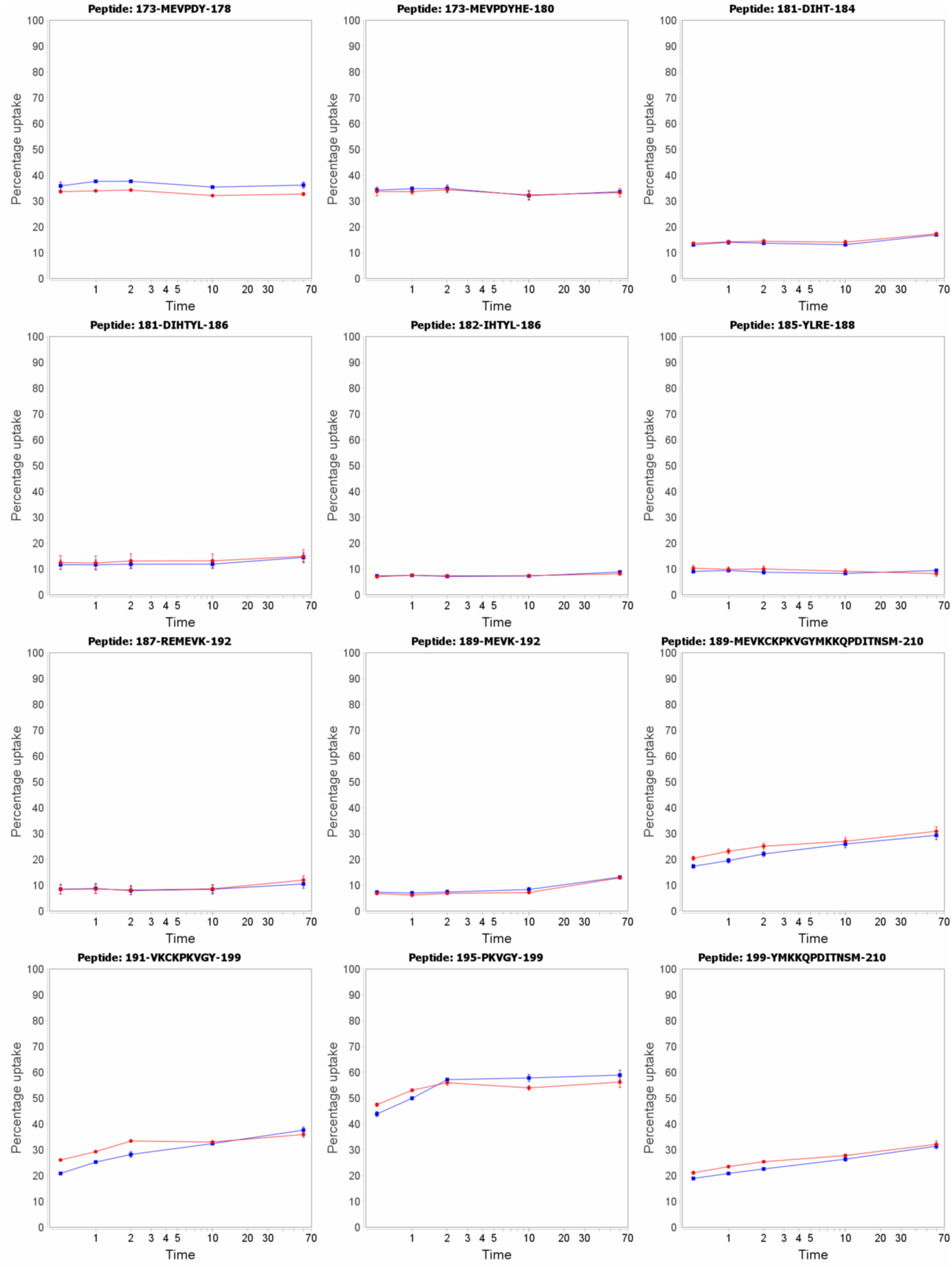
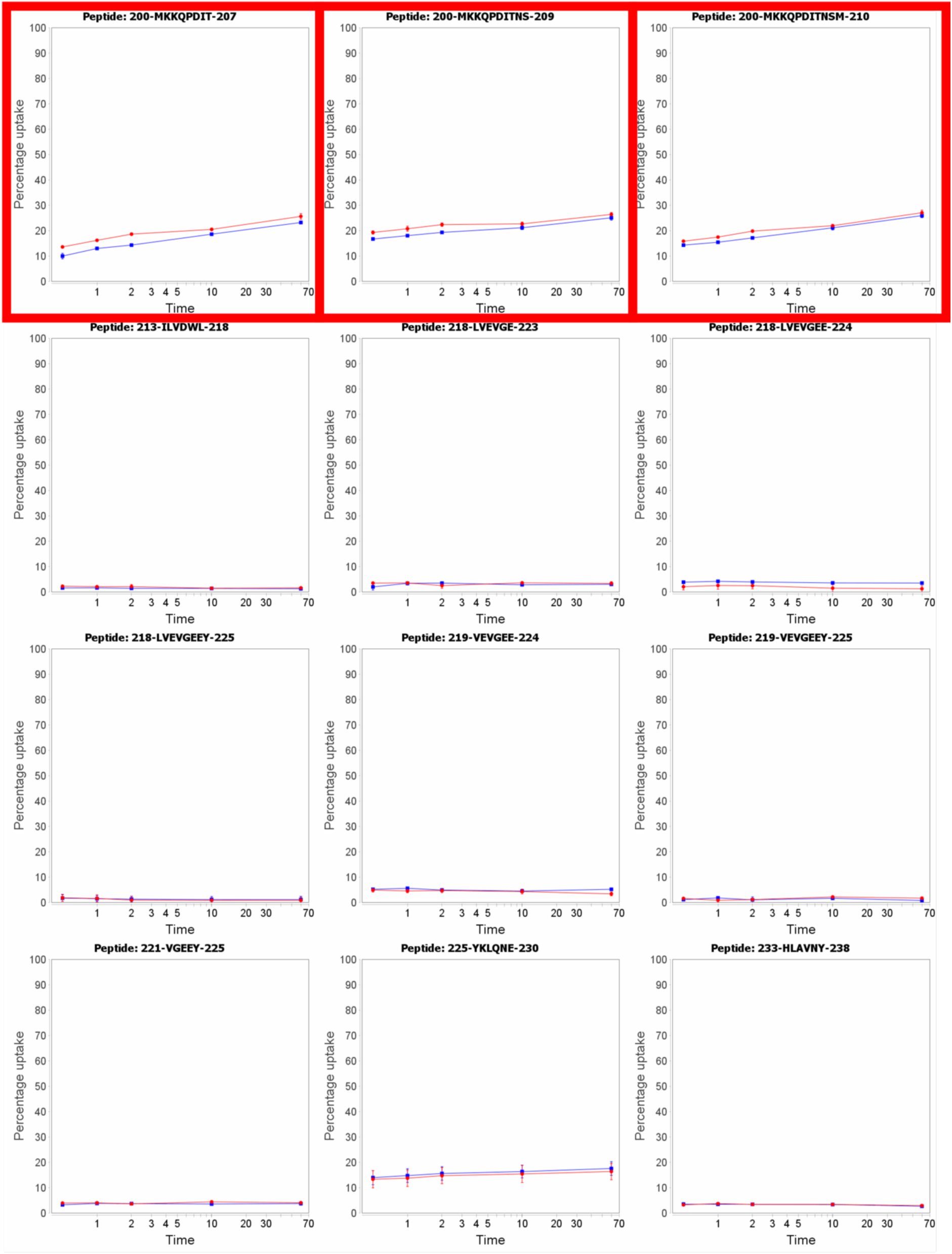
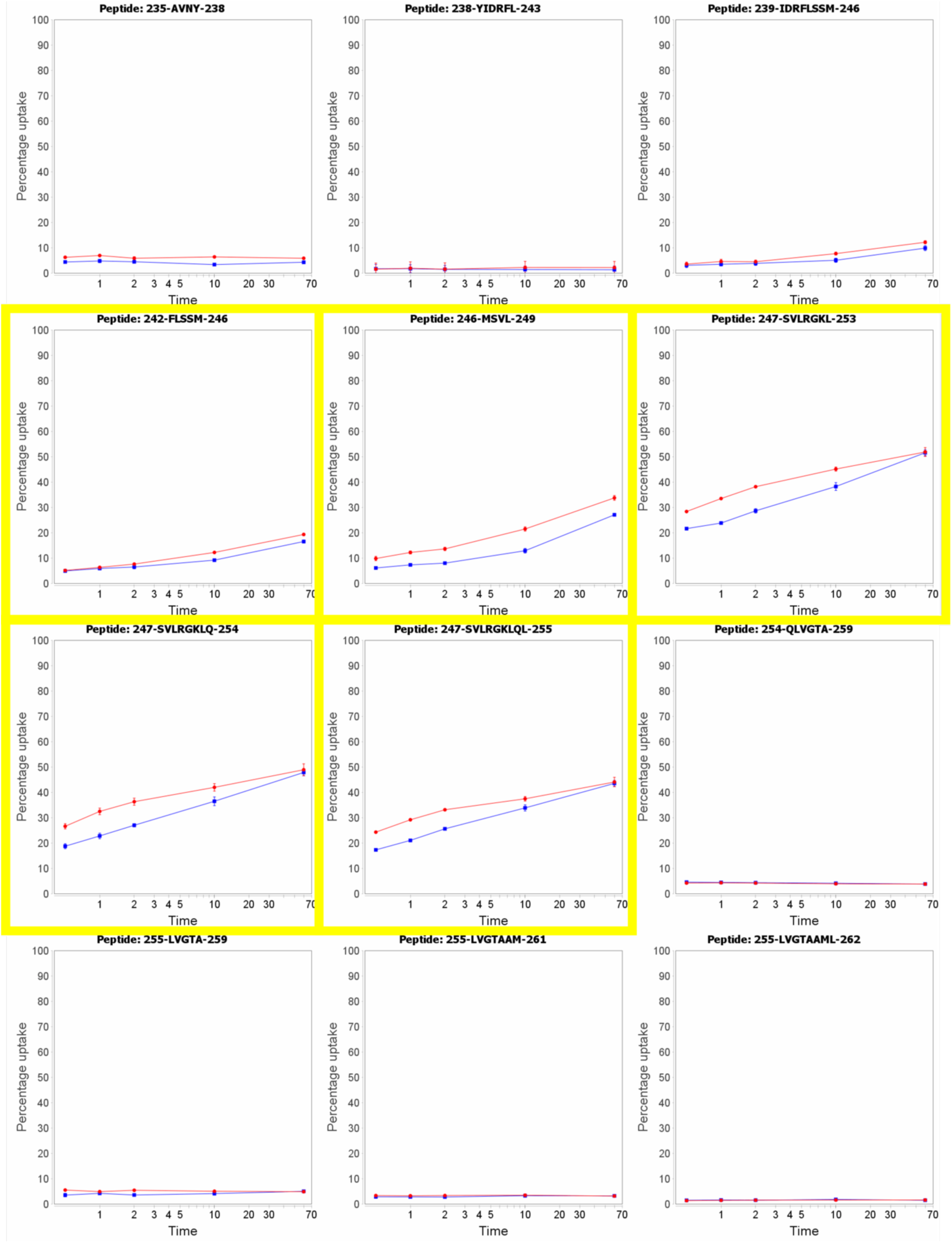
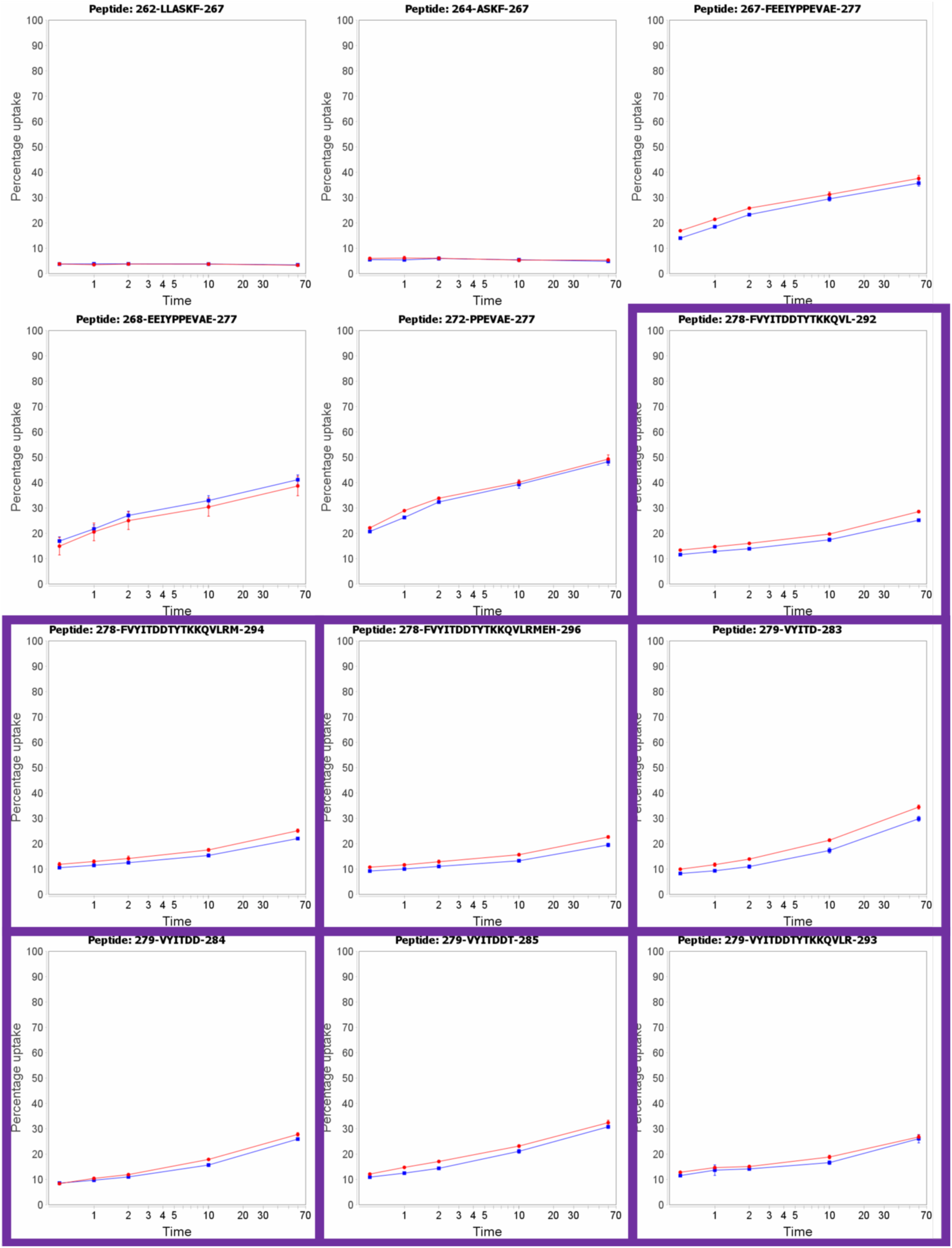
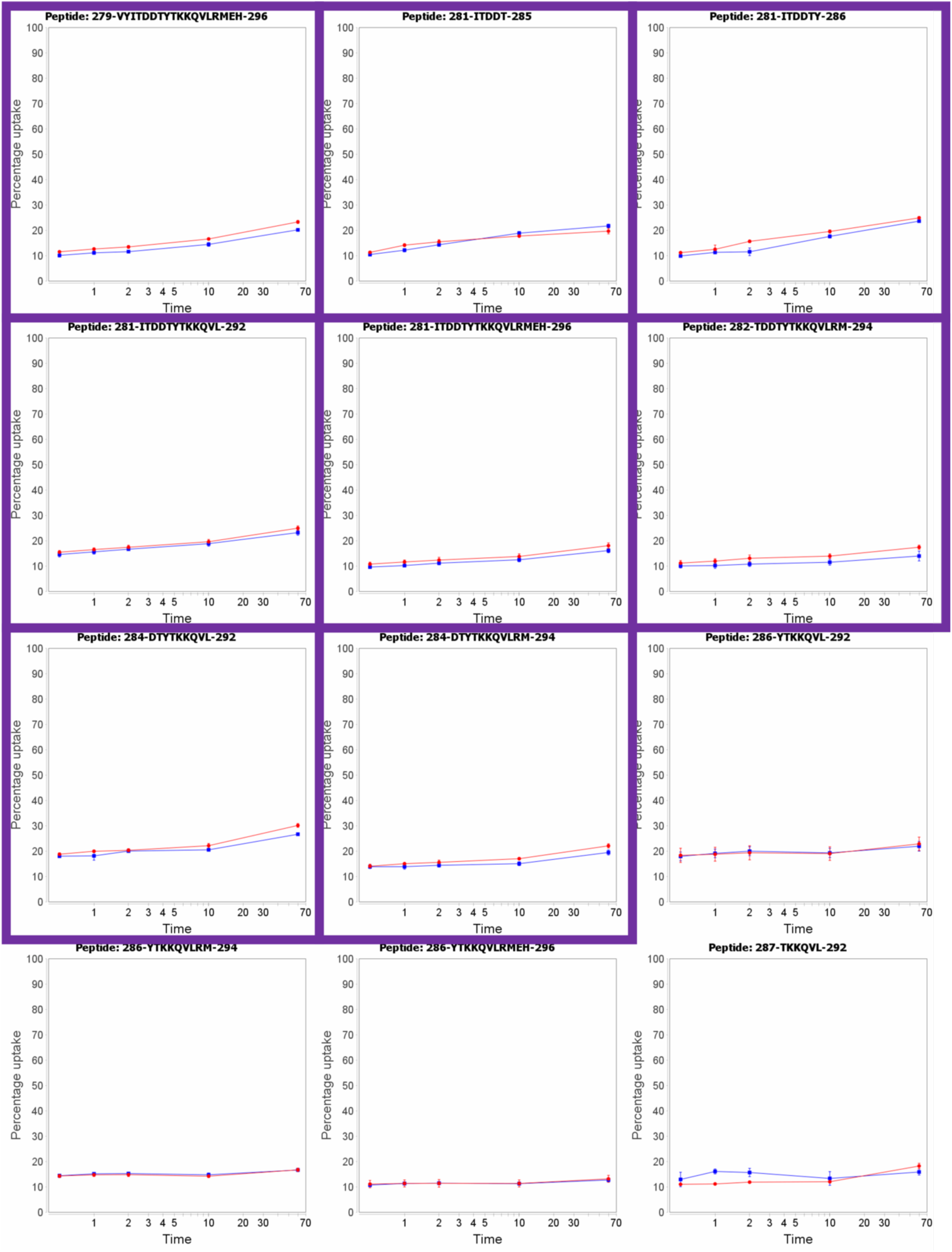
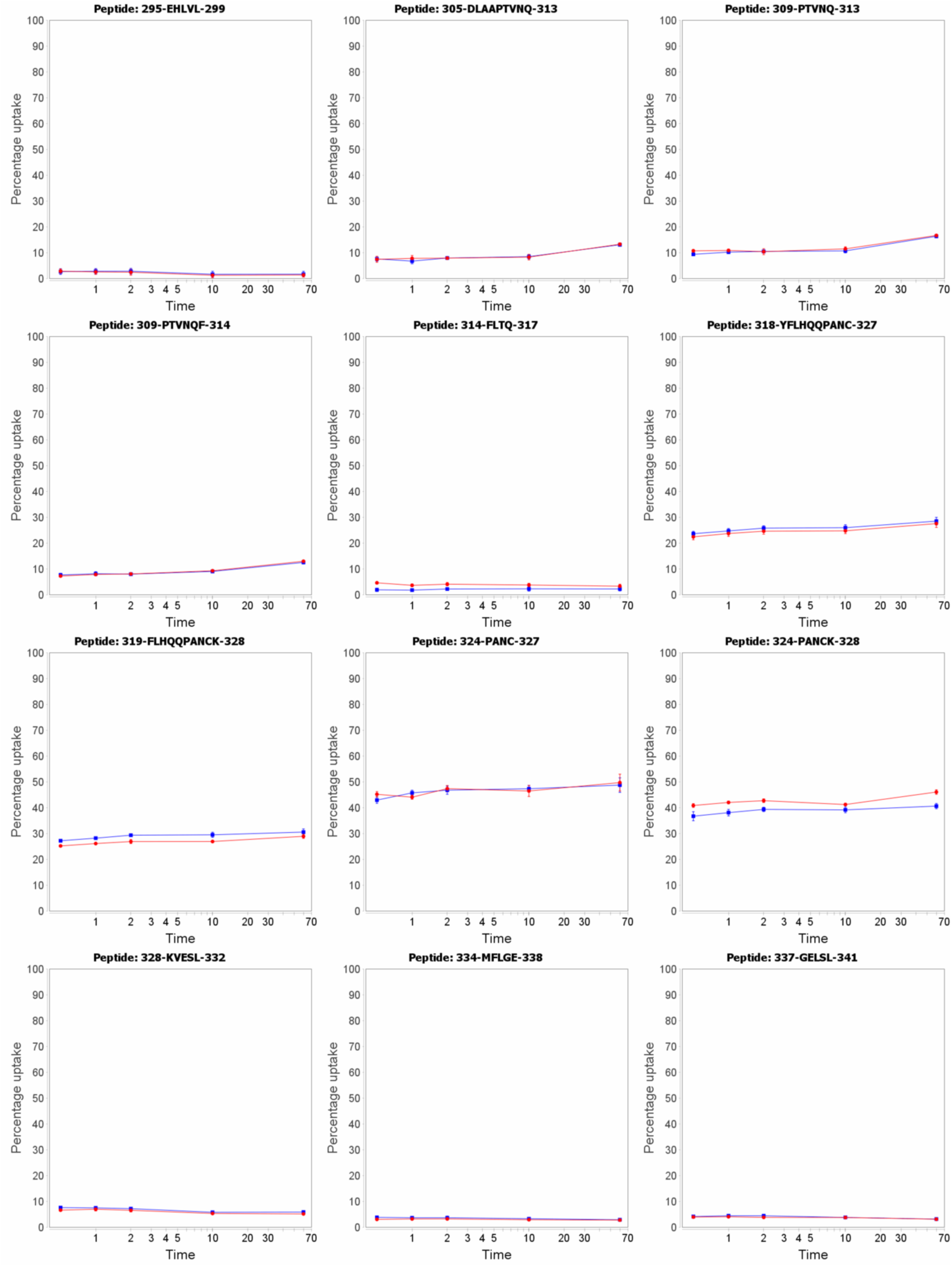
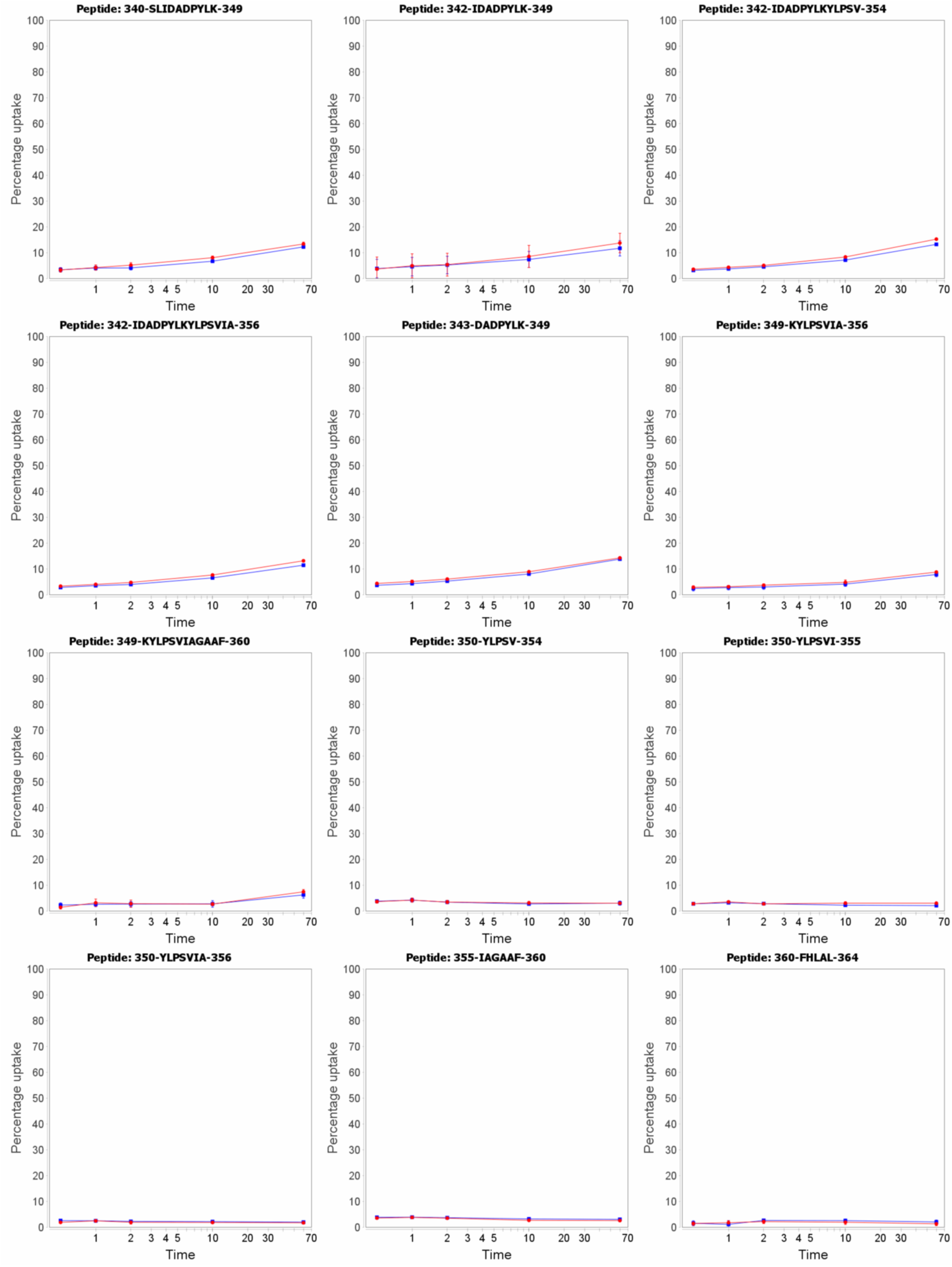
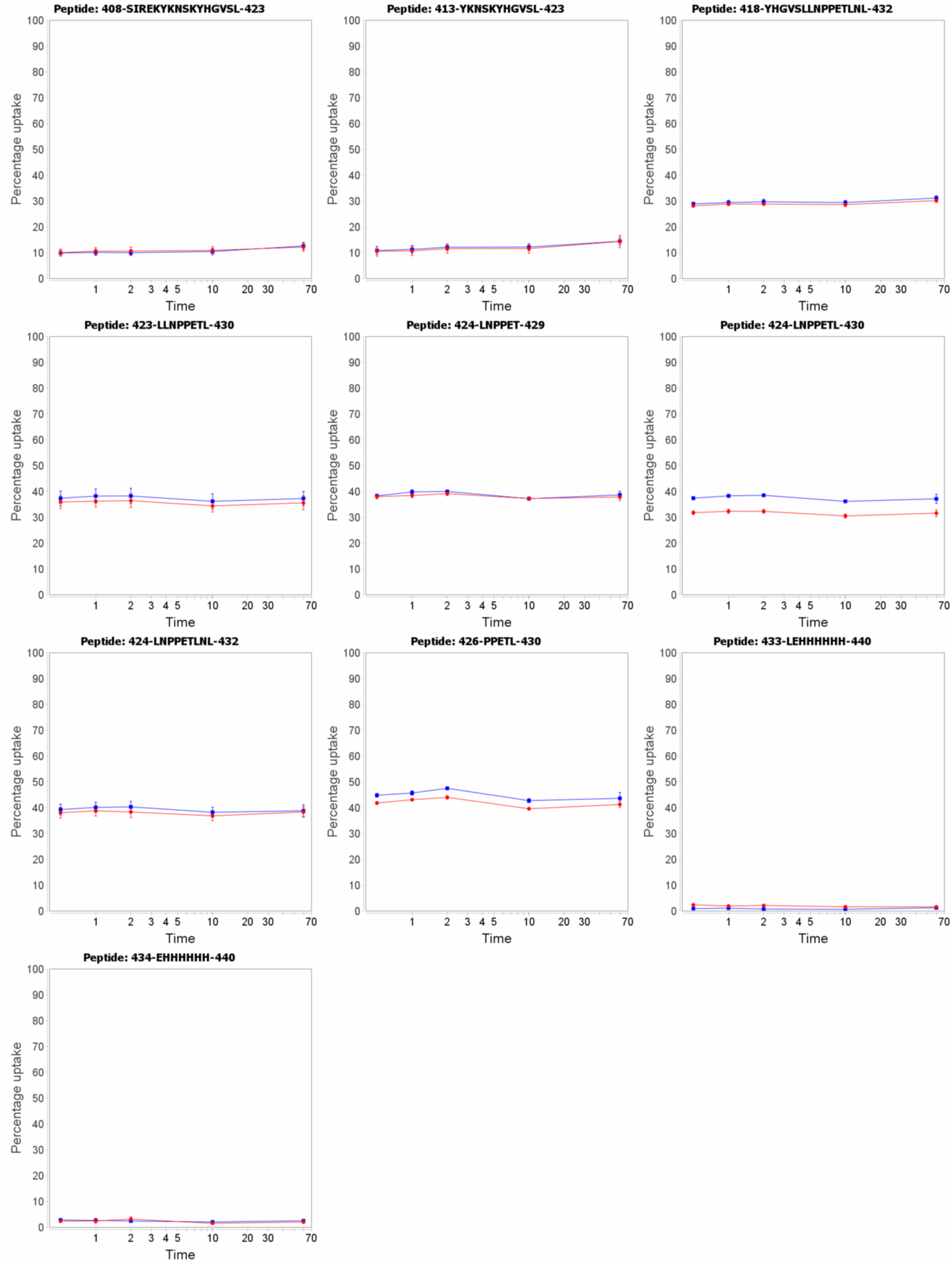
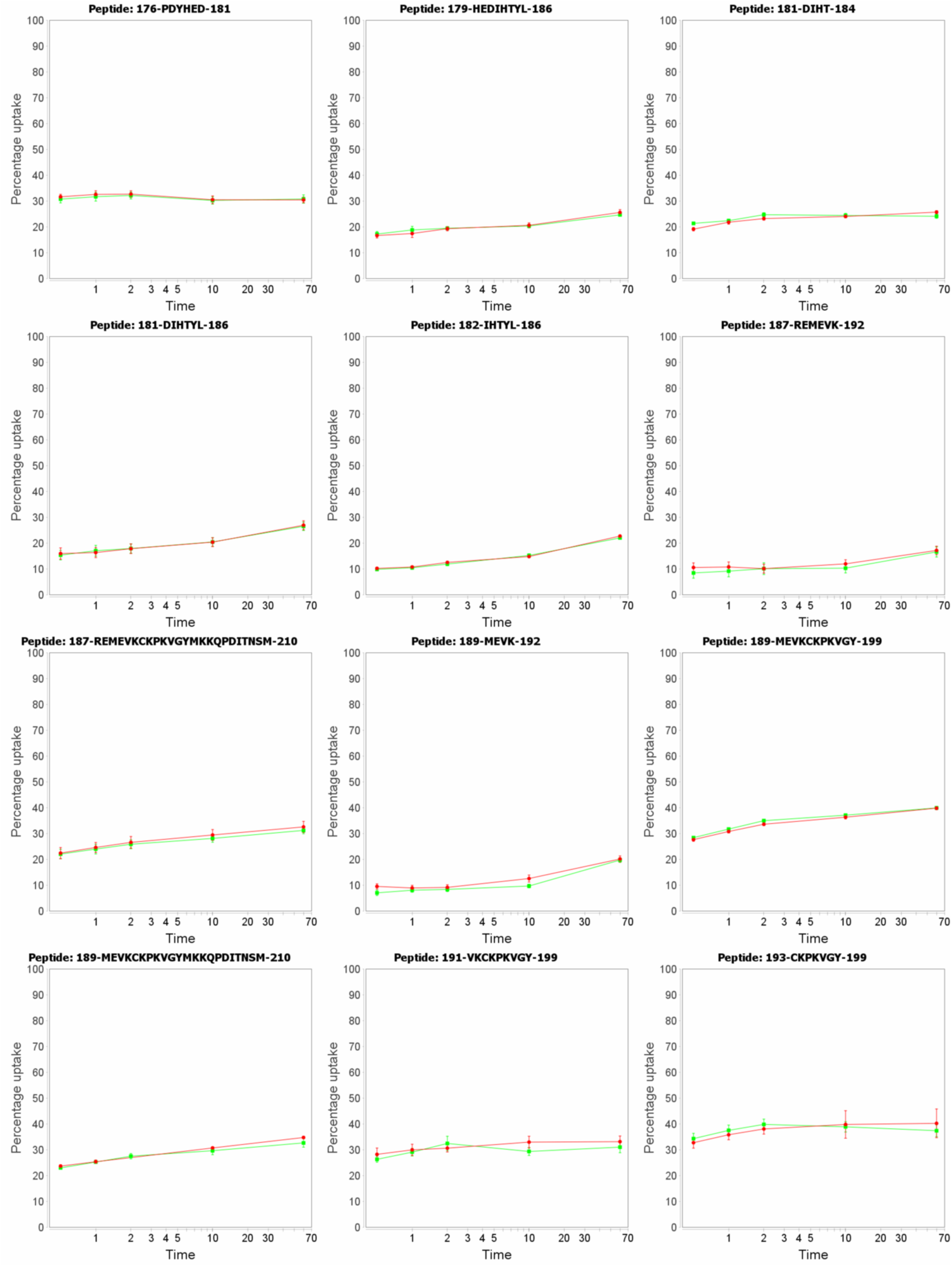
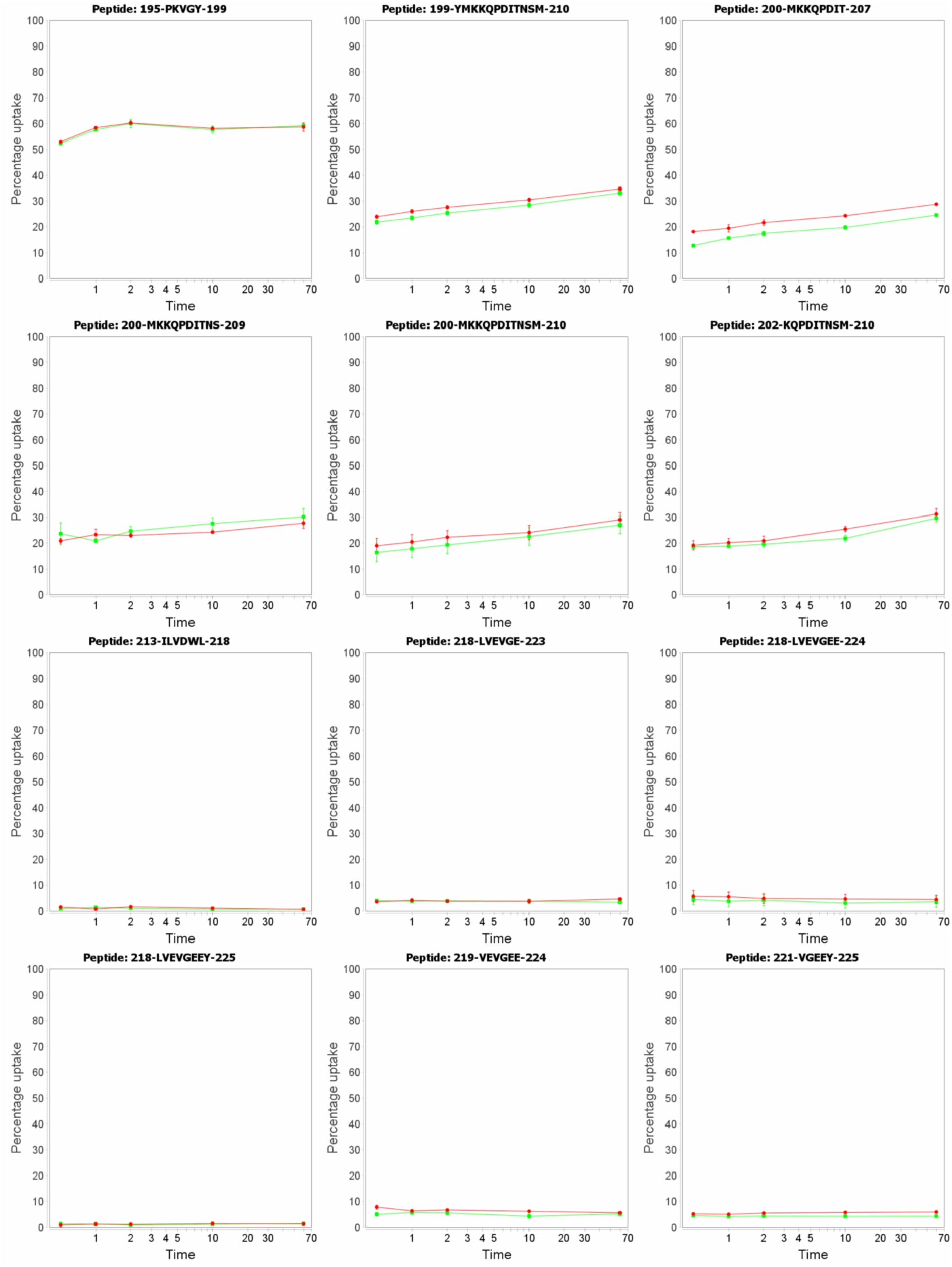
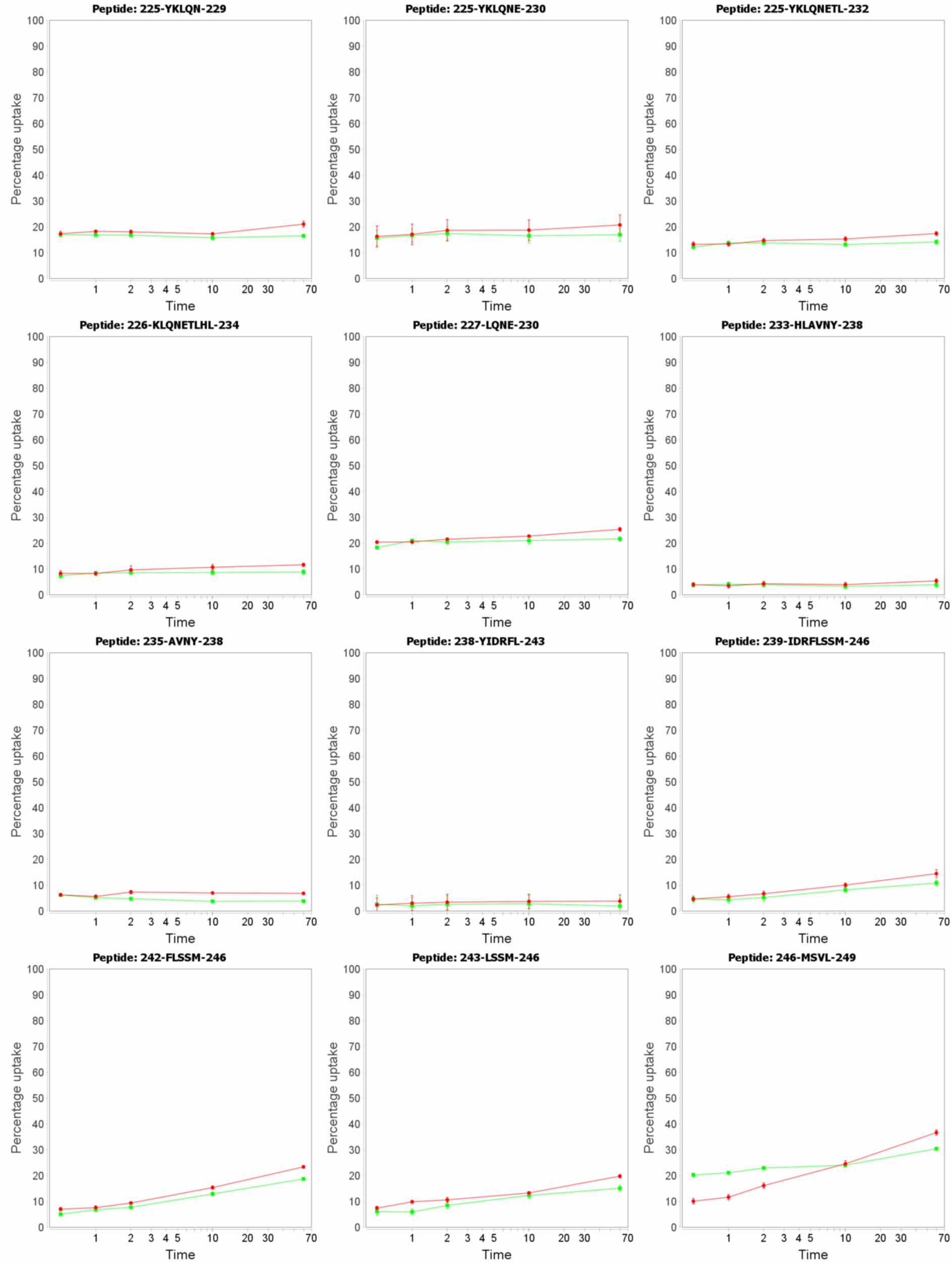
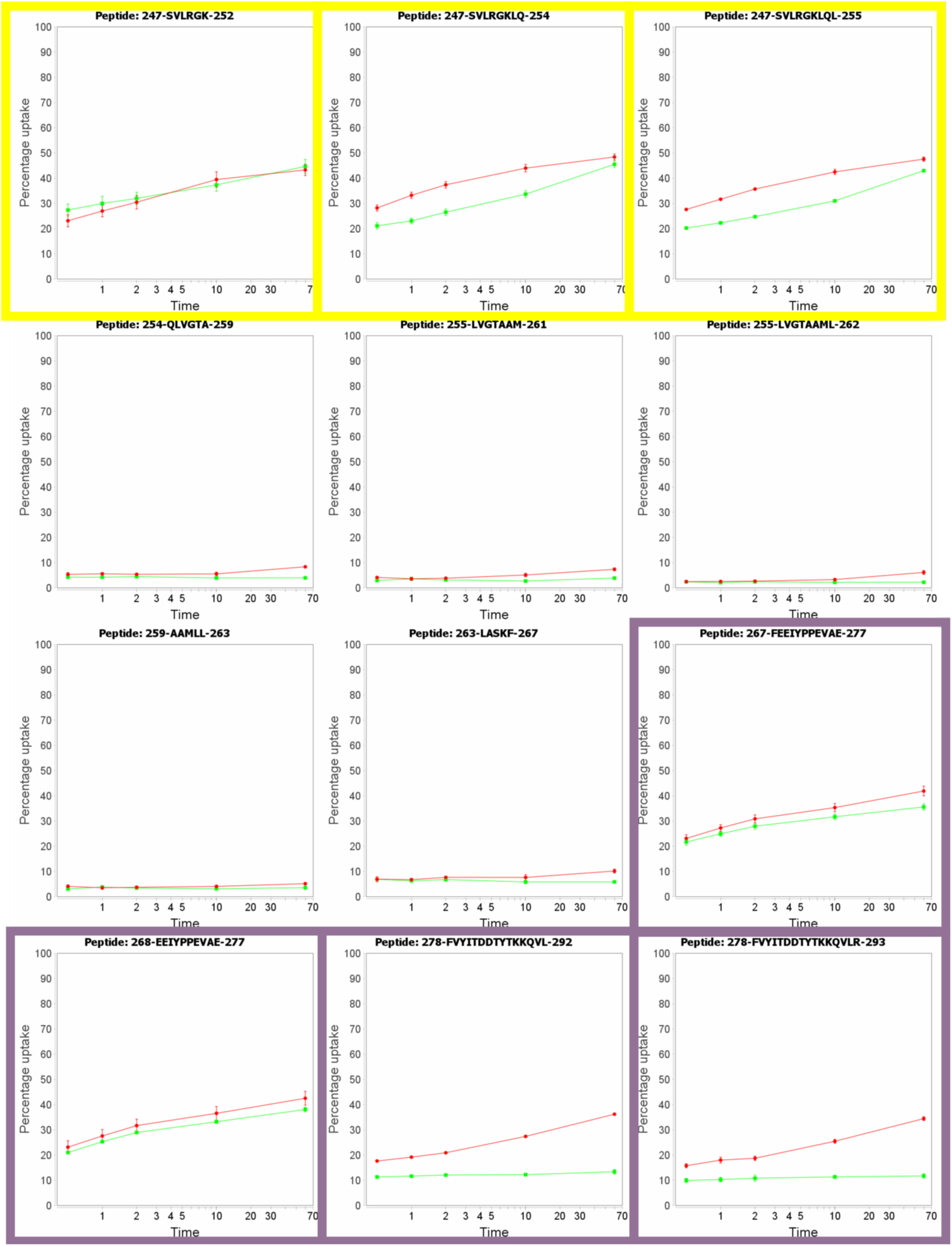
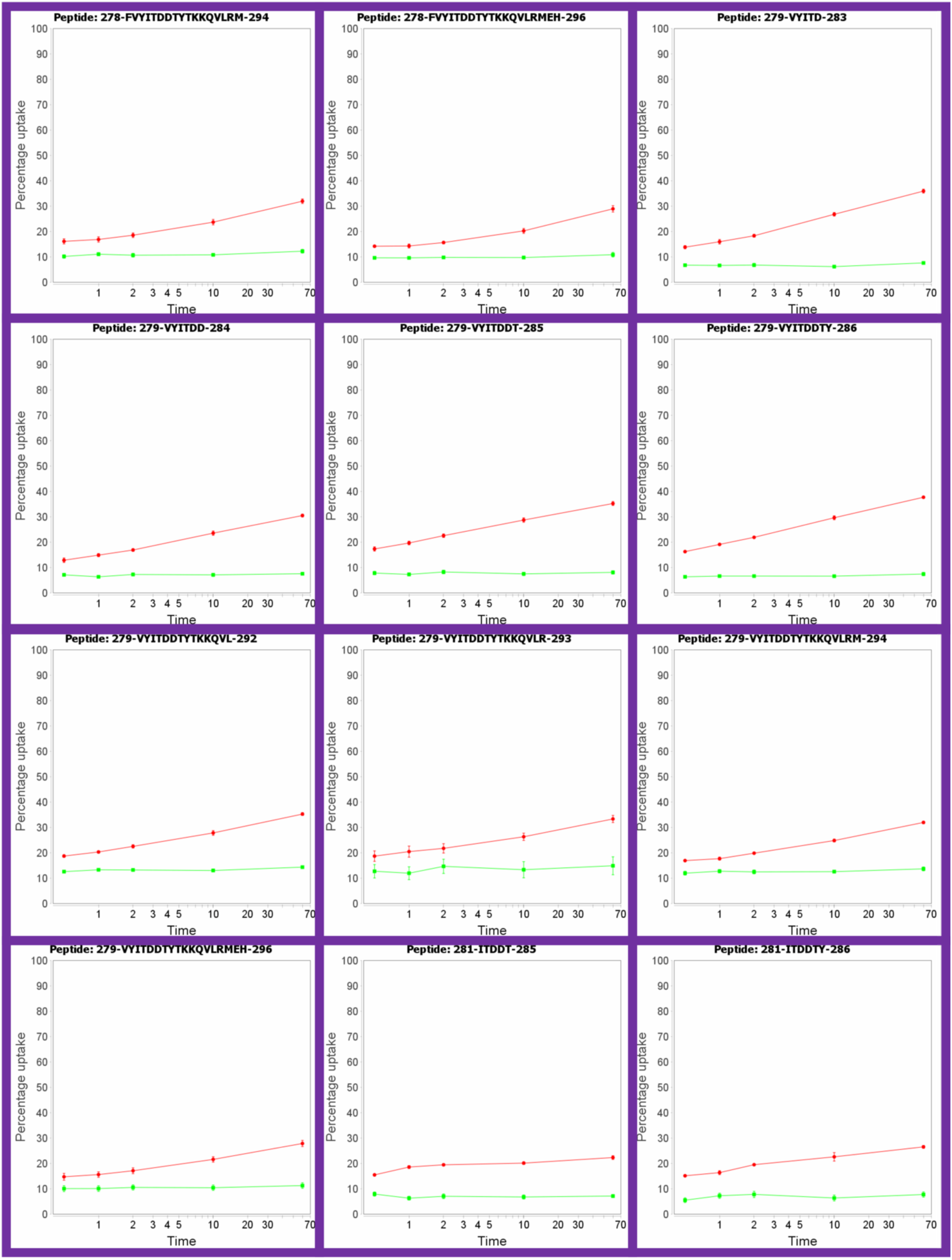
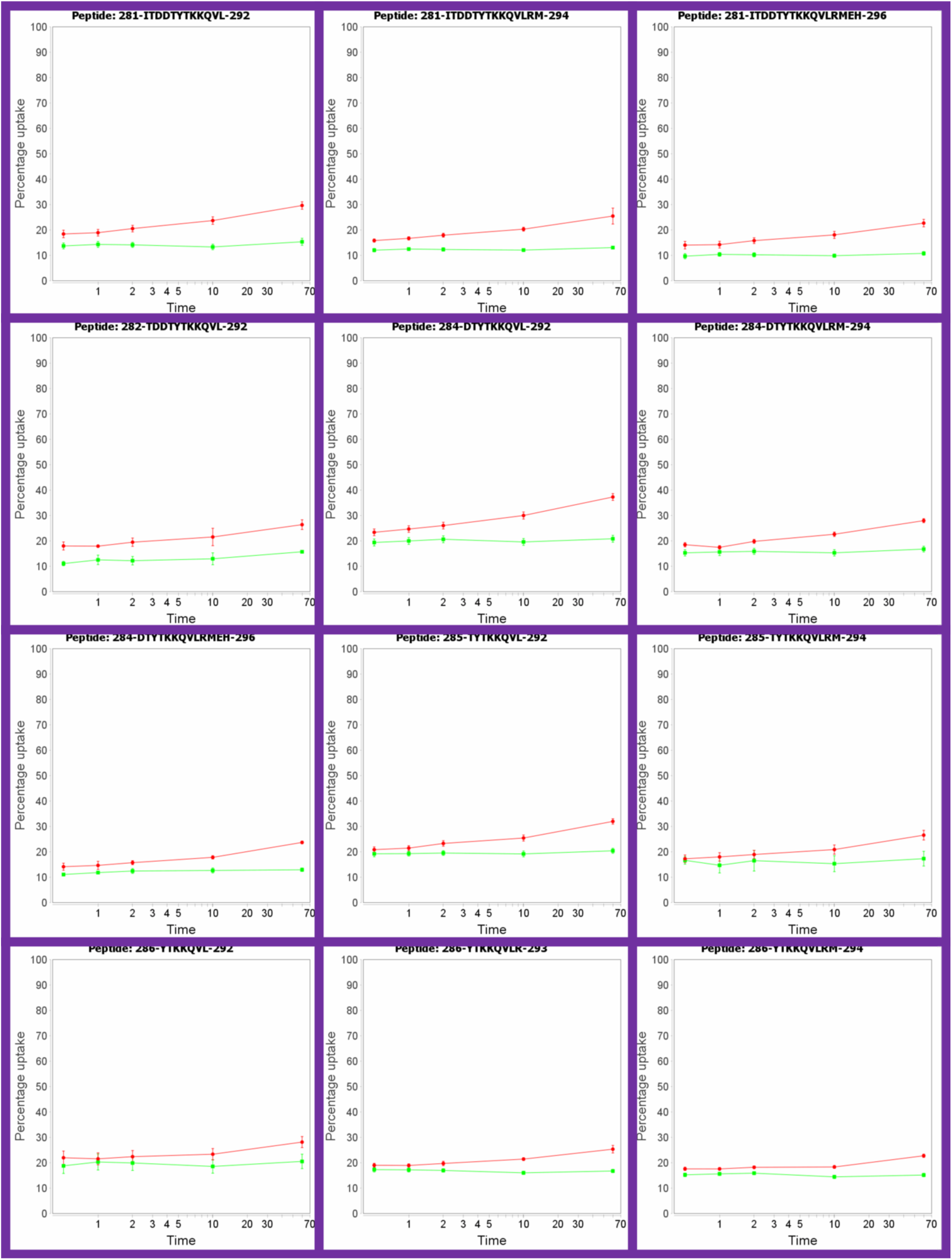
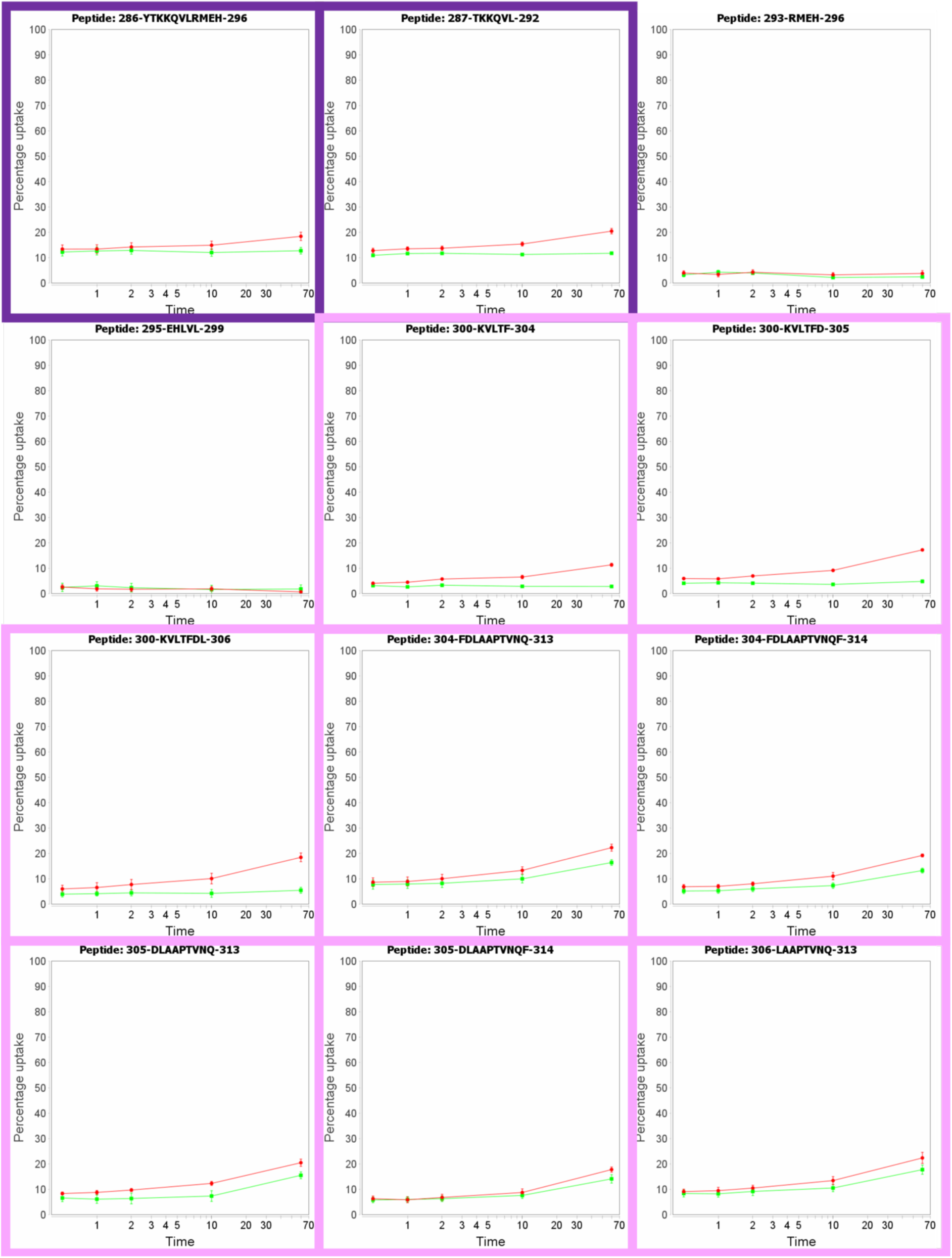
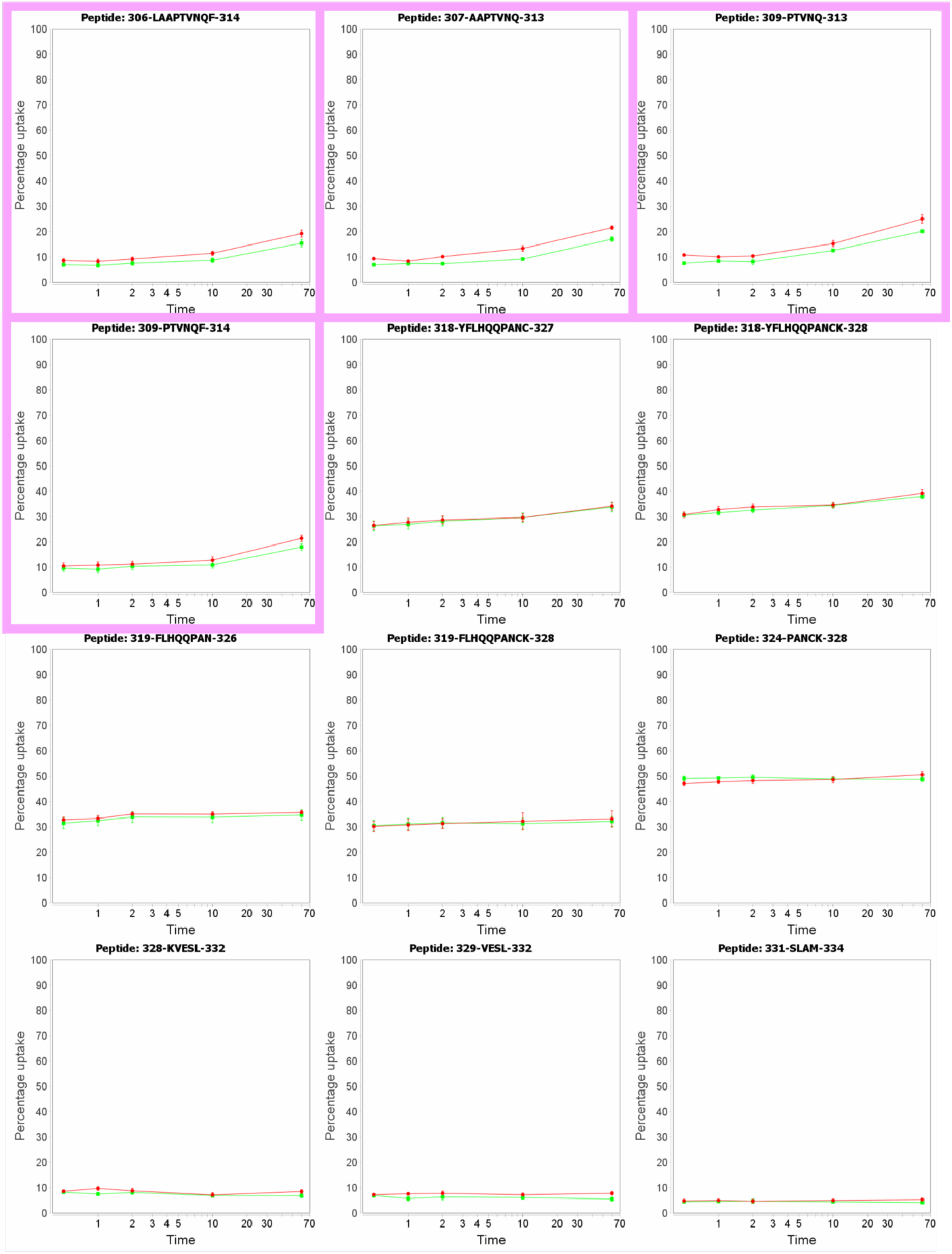
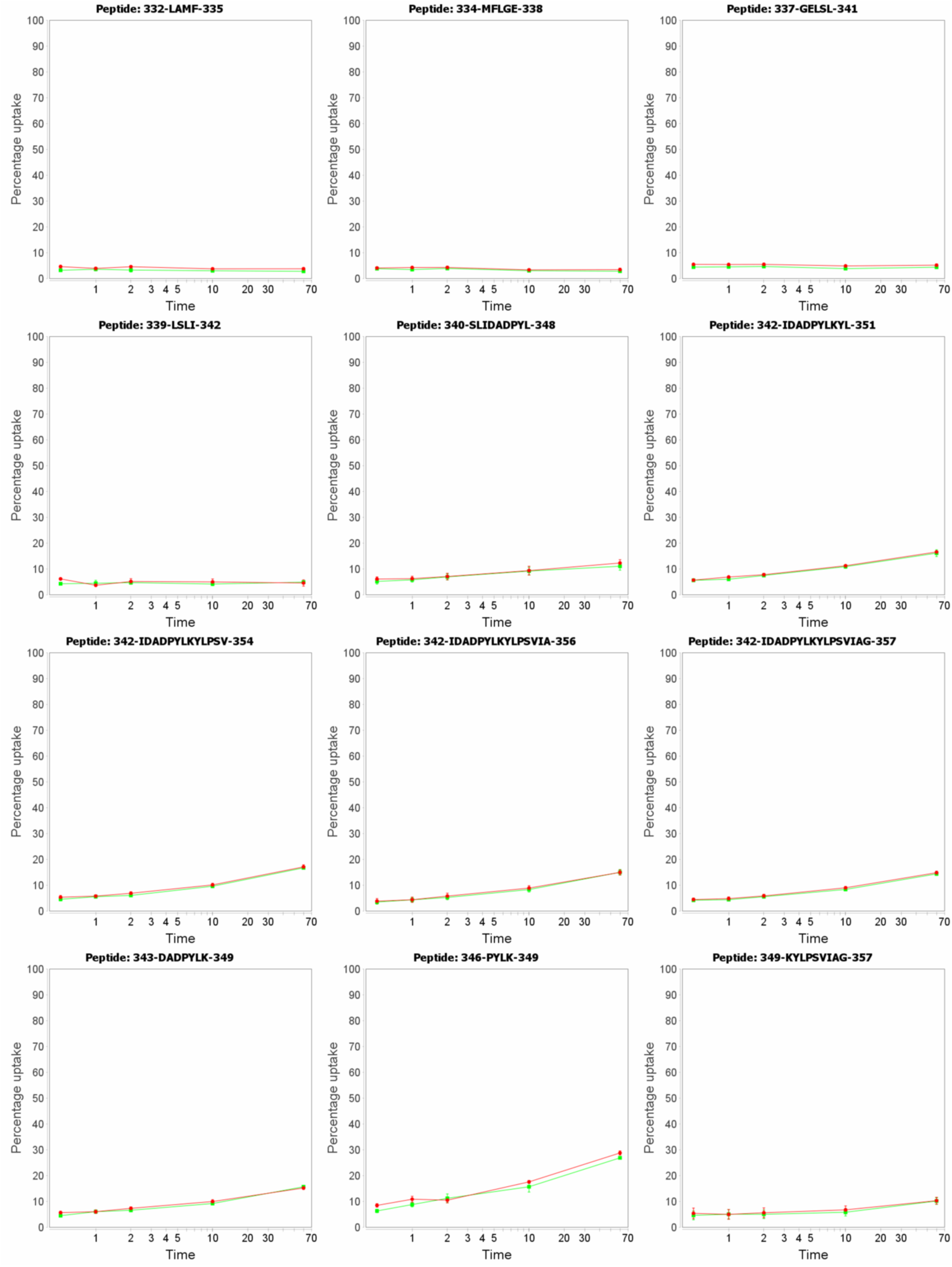
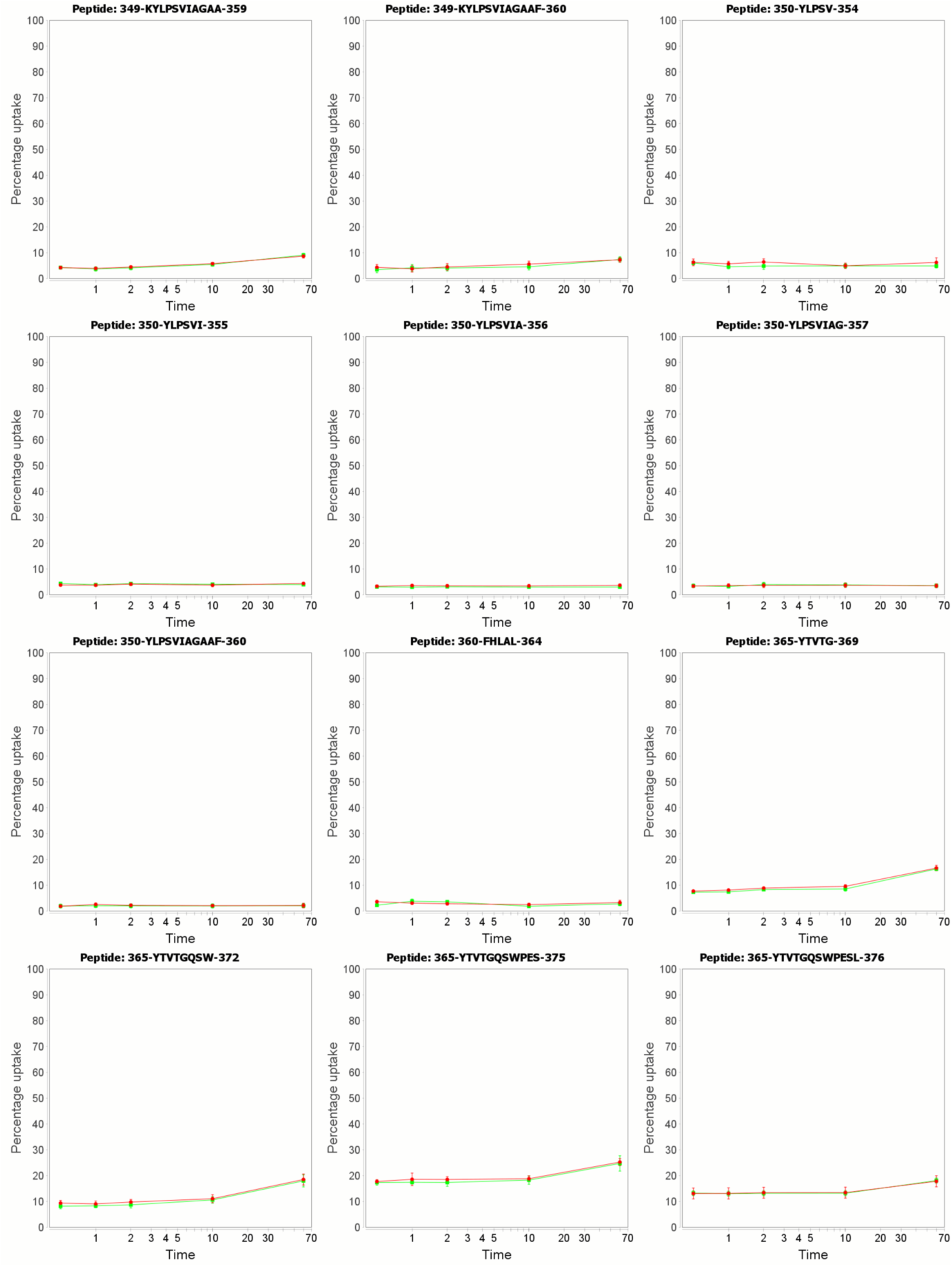
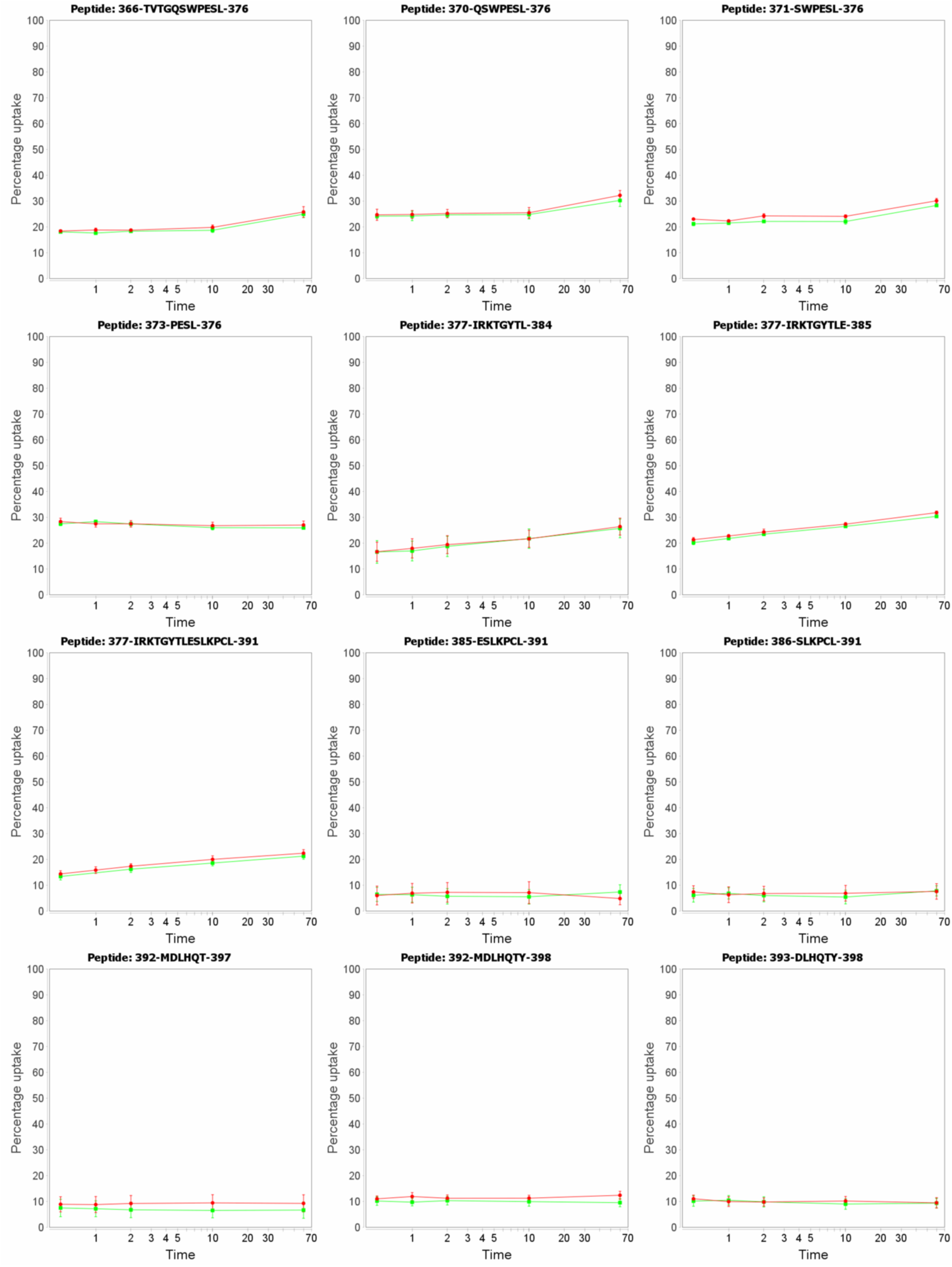
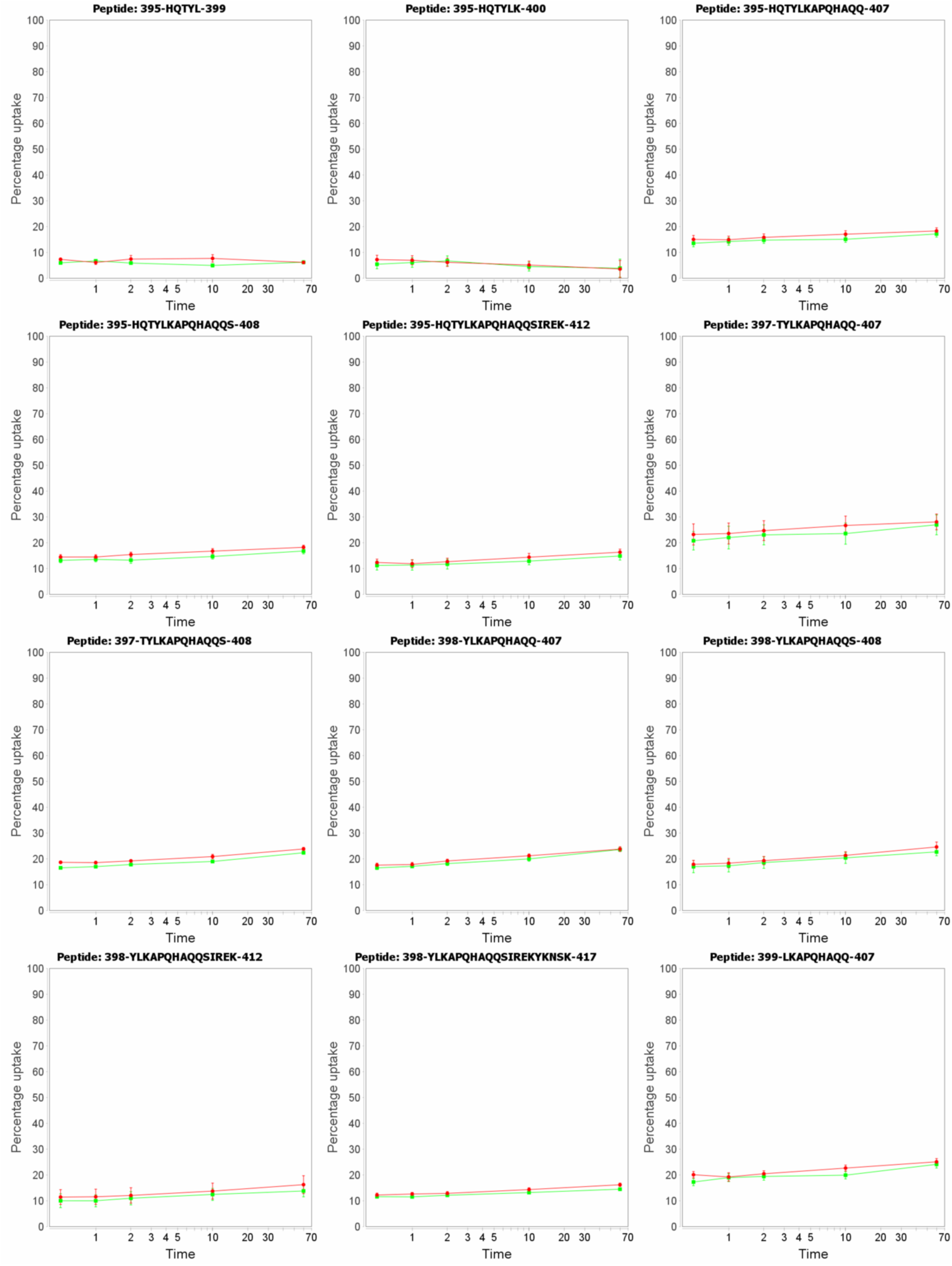
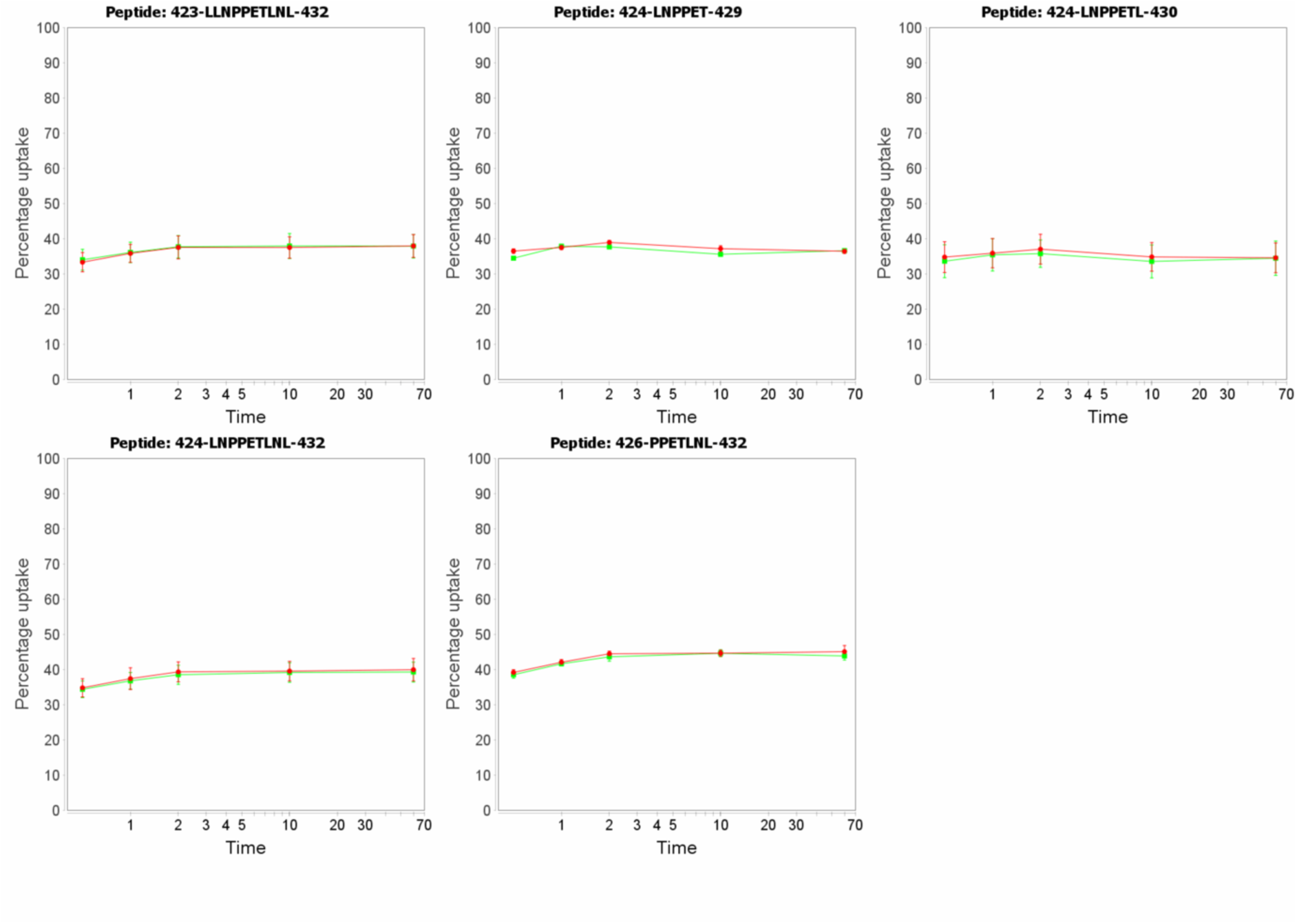
Hydrogen-deuterium exchange mass spectrometry to identify the cyclin A SKP2 binding site. Related to Figure 2. Uptake plots for cyclin A peptides in the absence (red) or presence of SKP1-Δ20SKP2N (blue) or p27KIP1M (red, green). Time in minutes. Highlighted peptides are boxed. Coloring as in Figure 2.

**Supplementary Figure S5.**
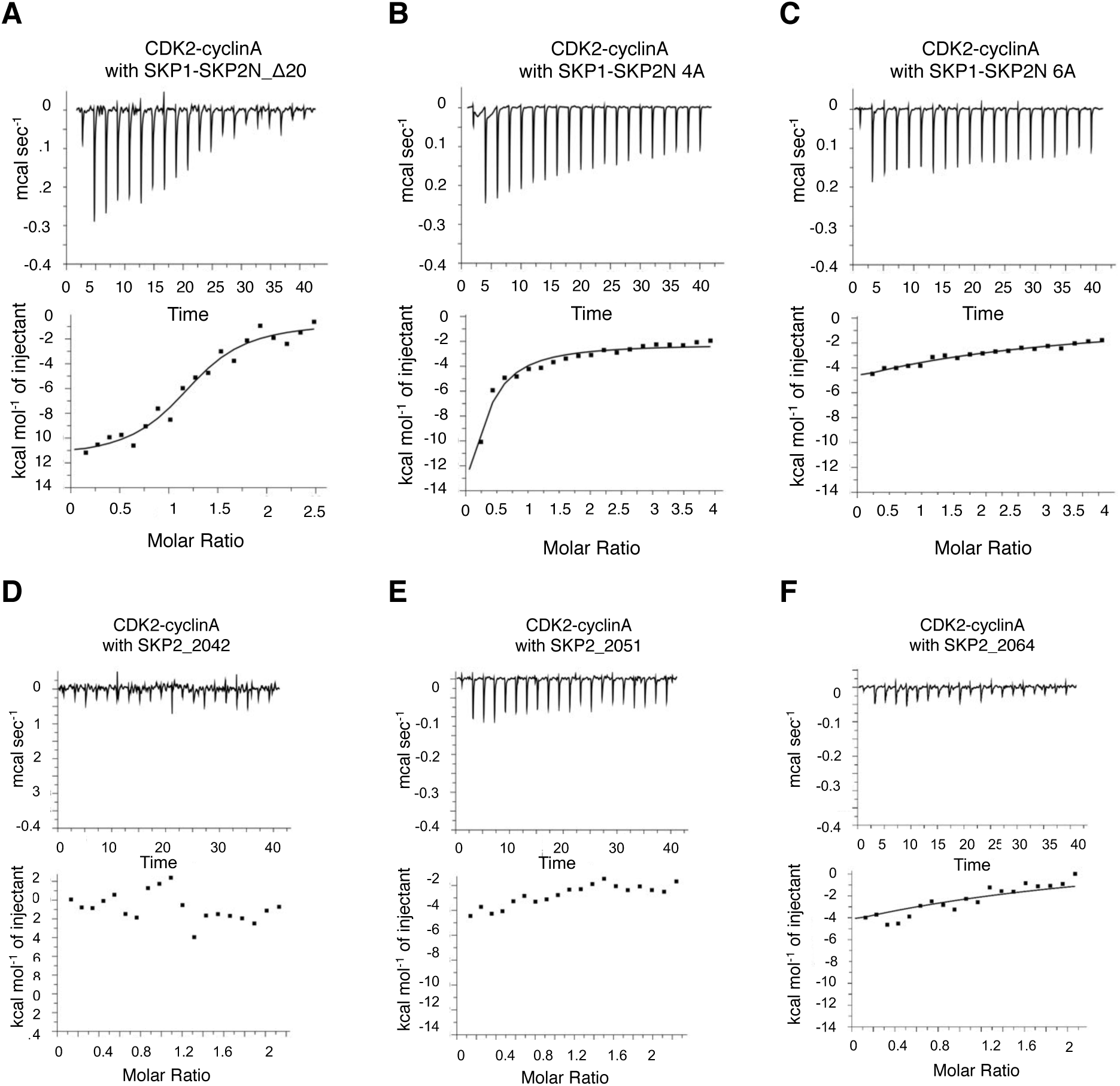
Characterization of the interaction between the SKP2 N-terminal sequence and cyclin A. Related to Table 2. (**A-C**) Isothermal titration calorimetry to characterize the interaction between CDK2-cyclin A and various mutant and truncated SKP2 constructs. (A) SKP1-SKP2N_Δ20, (B) SKP1-SKP2N_4A (L32A, L33A, S39A, L41A), and (C) SKP1-SKP2N_6A (W22A, W24A, L32A, L33A, S39A, L41A). (**D-F**) N-terminal SKP2 peptides 20-42 (D), 20-51, (E) and 20-64 (F) do not bind to cyclin A in the absence of the F-box bound *in trans* to SKP1. The isothermal titration calorimetry experiments were repeated twice using independently prepared protein samples.

**Supplementary Figure S6.**
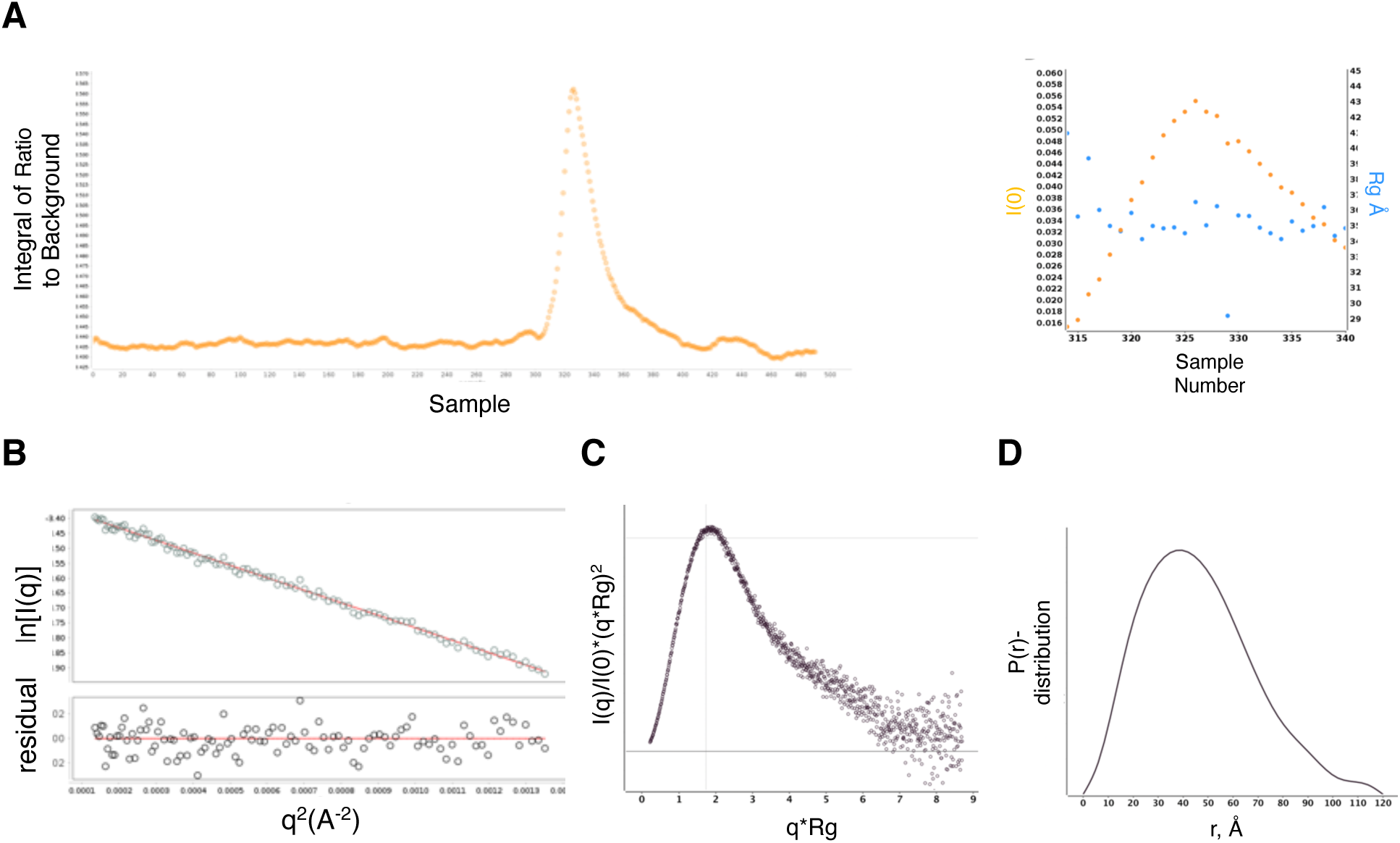
SAXS analysis of a CDK2-cyclin A-SKP1-SKP2 complex. Related to Figure 2. Supporting experimental plots for analysis of a SKP1(Δ38-43, Δ69-83)-SKP2(20-140-Δ60-74)-CDK2-cyclin A complex by SAXS. (**A**) FPLC size-exclusion chromatogram and fraction analysis. (**B**) Guinier plot. The sample monodispersity and absence of aggregation was assessed by a linear fit of the Guinier plot (ln(I) vs. q2) (**C**) Kratky plot. The dimensionless Kratky plot (q2 x I(q)/I(0)) suggests a globular and relatively compact particle (maximum at qRg of 1.73 with a height of 1.1), with some indications of a multi-domain characteristic with only limited flexibility overall (indicated by the drop of the curve between qRg of 4 – 6) (Perry and Tainer, 2013). (**D**) Pairwise distance distribution. In agreement, the pairwise distance distribution p(r) showed a bell shape indicating a fairly globular particle. A shoulder at d = 80 – 100 Å indicated that a domain sticks out from the globular shape, whereas the tail at d = 105-120 Å suggested the presence of a short flexible linker protruding from the particle.

**Supplementary Figure S7.**
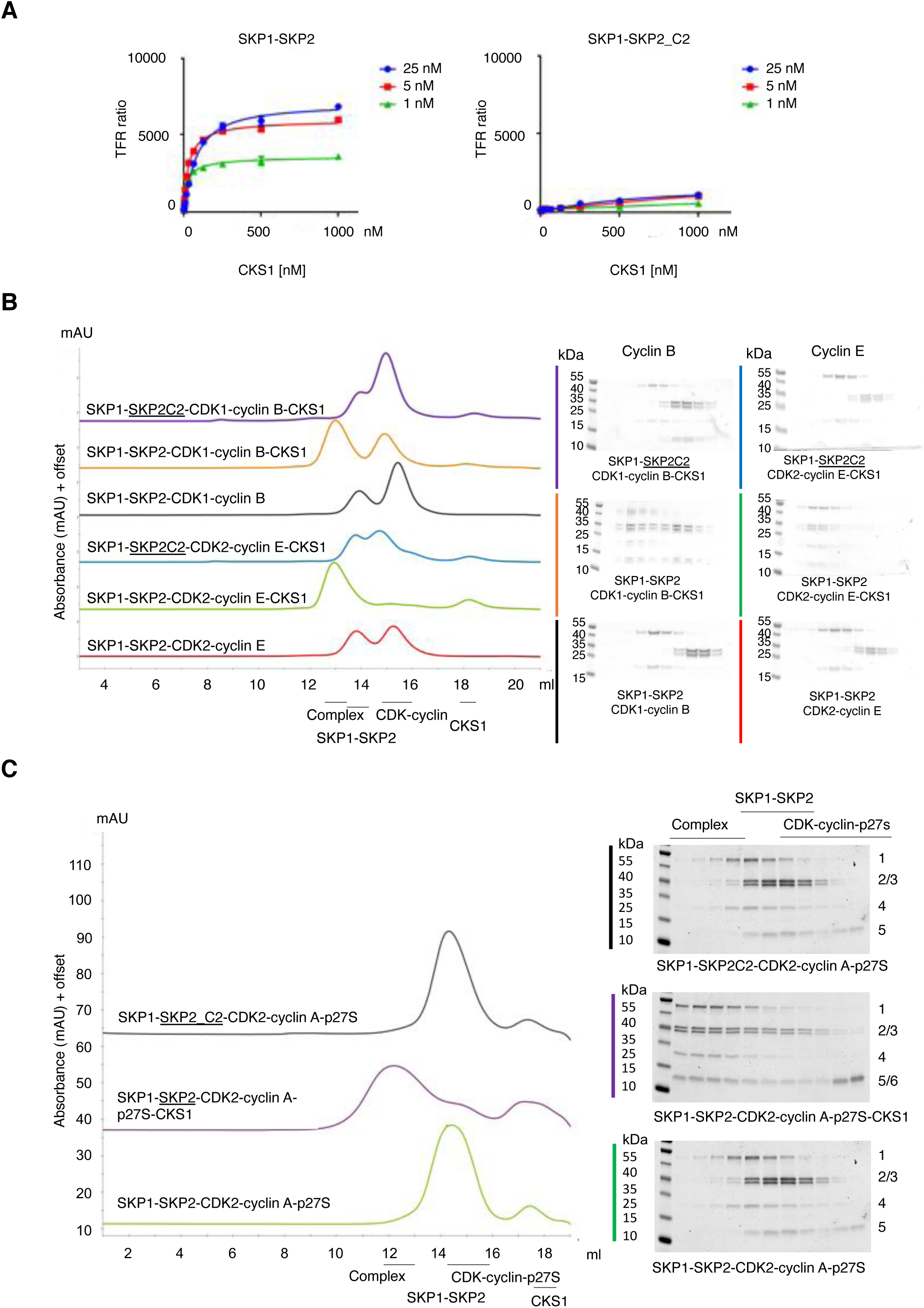
Characterization of the interaction between SKP2 and CKS1. Related to Figure 3. (**A**) Homogenous-time-resolved fluorescence (HTRF) assay to measure the binding of CKS1 to SKP1-SKP2 (left hand panel) and SKP1-SKP2_C2 (right hand panel). SKP2 (residues 101-436) co-expressed with SKP1 (1-163, Δ38-43 and Δ69-81)), and CKS1 (residues 5-73) are N-terminally tagged with GST or the AviTag respectively. The derived K_d_ values were dependent on the SKP1-SKP2 concentration (at 25, 5 and 1 nM, equal to 81.0, 31.3 and 20.8 nM respectively) from which it can be deduced that the actual K_d_ value for the interaction is circa < 1 nM (the lowest SKP1-SKP2 concentration used in the experiment). Replacement of SKP2 F393 with a G-S pair in the construct SKP1-SKP2_C2 severely impairs CKS1 binding to SKP2. The error bars represent the SEM. At least two independent repeats were carried out for each experiment. (**B**) The CDK2-CKS1 or CDK1-CKS1 interface is sufficient to maintain a stable pentameric complex in the absence of a cyclin-SKP2 interaction. Analytical size exclusion chromatography (SEC) assesses the ability of CDK1-cyclin B, CDK2-cyclin A and CDK2-cyclin E to form complexes with SKP1-SKP2 in the presence and absence of CKS1. SKP1, SKP2 and CKS1 are all full-length. Chromatograms represent single experiments and are displayed offset along the y-axis for clarity. (**C**) The interaction of p27KIP1 with SKP2 is dependent on CKS1. Analytical SEC shows that p27KIP1 bound to CDK2-cyclin A forms a complex with SKP2 in the presence of CKS1 (compare magenta and green traces). When the CKS1 binding site on SKP2 is compromised (by introduction of the SKP2_C2 mutation) CDK2-cyclin A-CKS1 and SKP1-SKP2 complexes are detected (black trace). Chromatograms are representative of 2 biological replicates carried out using either p27KIP1M (residues 1-106) or p27KIP1S (residues 23-106). Experiment using p27KIP1S is shown and chromatograms are displayed offset along the y-axis for clarity. SKP2, CDK2, cyclin A, SKP1, p27KIP1S and CKS1 are identified by numbers 1-6 respectively. CKS1 (9.6 kDa) and p27KIP1S (9.9 kDa) co-migrate.

**Supplementary Table S1.** Data collection and refinement statistics. Related to Figure 2.

|  | CDK2-cyclin Amut5 |
| --- | --- |
| **Data collection** |  |
| Space group | P12<sub>1</sub>1 |
| Cell dimensions |  |
| *a*, *b*, *c* (Å) | 40.3 137.8 109.7 |
| $\alpha$, $\beta$, $\gamma$ (°) | 90.0 99.8 90.0 |
| Resolution (Å) | 85.10 - 2.43 (2.52 - 2.43) |
| *R*<sub>sym</sub>/*R*<sub>merge</sub> | 0.067 (0.587) |
| *I*/σ*I* | 12.0 (1.3) |
| Completeness (%) | 99.6 (98.7) |
| Redundancy | 1.9 (1.9) |
| **Refinement** |  |
| Resolution (Å) | 85.10 - 2.43 (2.52 - 2.43) |
| No. reflections | 44272 / 2129 |
| *R*<sub>work</sub> / *R*<sub>free</sub> (%) | 20 / 25.1 |
| No. atoms |  |
| Protein | 8366 |
| Ligand/ion | 22 |
| Water | 97 |
| *B*-factors |  |
| Protein | 47.2 |
| Ligand/ion | 51.5 |
| Water | 32.1 |
| R.m.s. deviations |  |
| Bond lengths (Å) | 0.005 |
| Bond angles (°) | 1.38 |

• Structure was determined from a single crystal following screening. • Values in parentheses are for highest-resolution shell.

**Supplementary Table S2.** Kinase activity of CDK2-cyclin A. complexes. Related to Figure 2. Complexes were assayed using a p107 substrate peptide using the ADP-Glo^TM^ assay format as described in the methods. ^1^ND (not determined). ^2^Experiments were repeated twice and error bars correspond to the range of values. ATP concentration, 75 μM. ^3^Experiments were performed in triplicate and error bars correspond to the range of values. ATP concentration, 25 μM.

|  | V<sub>max</sub> | K<sub>m</sub> | k<sub>cat</sub> (1/s) | k<sub>cat</sub>/K<sub>m</sub><br>(nM/s) |
| --- | --- | --- | --- | --- |
| <sup>2</sup>CDK2-cyclin A | 5.9 ± 1.2 | 40.7 ± 17.3 | 2.4 | 0.1 |
| <sup>2</sup>CDK2-cyclin A-SKP1-SKP2 | 7.6 ± 0.5 | 12.3 ± 2.5 | 3.1 | 0.2 |
| <sup>2</sup>CDK2-cyclin A-SKP1-SKP2<sup>S64A</sup> | 6.8 ± 0.7 | 15.1 ± 4.3 | 2.7 | 0.2 |
| <sup>2</sup>SKP1-SKP2<sup>S64A</sup> | <sup>1</sup> ND | ND | ND | ND |
| <sup>3</sup>CDK2-cyclin A | 19.8 ± 3.4 | 58.66 ± 19.4 | 7.9 | 0.1 |
| <sup>3</sup>CDK2-cyclin A-p27KIP1 | ND | ND | ND | ND |
| <sup>3</sup>CDK2-cyclin Amut5 | 19.5 ± 2.1 | 34.01 ± 8.5 | 7.8 | 0.2 |
| <sup>3</sup>CDK2-cyclin Amut7 | 21.3 ± 1.2 | 31.62 ± 4.3 | 8.5 | 0.3 |

**Supplementary Table S3.** SAXS data collection and processing. Related to Figure 2.

|  |  |
| --- | --- |
| **Data collection** |  |
| Beamline | BL21-Diamond |
| X-ray wavelength (KeV) | 12.4 |
| q range (Å) | 0.004-0.4420 |
| Concentration range (mg.mL<sup>-1</sup>) | 5 – 0.2 |
| Temperature (°C) | 10 |
| **Data Analysis** |  |
| I<sub>(0)</sub> | 3.88E-2 |
| Guinier q-region | 1-1383 |
| R<sub>g</sub> (Å) from Guinier (±SE) | 35.32 (± 0.39) |
| R<sub>g</sub> (Å) from GNOM (±SE) | 34.86 (± 0.55) |
| D<sub>max</sub> (Å) | 110 |
| Resolution (Å) | 16.1 |
| Oligomeric state | monomer |
| **Ab Initio modelling** |  |
| Number of models | 12 |
| **Software employed** |  |
| Primary data reduction | Scatter3 |
| Theoretical data fitting | Scatter3 |
| Envelope modelling | DAMMIF |
| Modelling | Coral and EOM |

